# Generative AI-driven artificial DNA design for enhancing inter-species gene activation and enzymatic degradation of PET

**DOI:** 10.1101/2025.05.08.652991

**Authors:** Manato Akiyama, Motohiko Tashiro, Ying Huang, Mika Uehara, Taiki Kanzaki, Mitsuhiro Itaya, Masakazu Kataoka, Kenji Miyamoto, Yasubumi Sakakibara

## Abstract

Conventional approaches to heterologous gene expression rely on codon optimization, which is limited to swapping synonymous codons and often fails to capture deeper adaptive changes. In contrast, naturally evolved orthologous genes between species often differ by more than just synonymous substitutions – they can include non-synonymous mutations, insertions, and deletions that confer functional adaptation to different host contexts. Here we present OrthologTransformer, a Transformer-based deep learning model that converts orthologous genes between species by learning from large-scale orthologous gene datasets curated for high-quality sequence alignments. The model recapitulates the full spectrum of evolutionary differences – from synonymous codon swaps to amino acid-changing mutations and indels – to predict a coding sequence optimized for a target species while preserving the protein’s function. In extensive validation across diverse bacterial species pairs, OrthologTransformer significantly increased the conversion accuracy of generated genes to native target sequences compared to the original source genes, even for pairs with stark disparities in GC content and optimal growth temperature. The Transformer’s context-aware designs also favored conservative amino acid substitutions, maintaining protein functional integrity. As a proof-of-concept, an OrthologTransformer-designed PETase gene for *Bacillus subtilis* from the host species *Ideonella sakaiensis* was synthesized and expressed, yielding robust PET plastic-degradation activity that surpassed synonymous codon-optimized controls. These results establish OrthologTransformer as a powerful tool for de novo cross-species gene adaptation, transcending the limits of traditional codon optimization and enabling more effective heterologous gene performance in synthetic biology applications.

## Introduction

Heterologous gene expression – expressing a gene in a non-native host organism – has become a cornerstone of modern biotechnology, enabling the production of recombinant proteins and enzymes in convenient expression systems. A fundamental challenge in this process is that different organisms exhibit distinct codon usage biases; the frequency with which synonymous codons are used can vary greatly between a gene’s native species and the host species^1,2^. To overcome this, codon optimization is routinely employed to modify a gene’s coding sequence without altering the amino acid sequence, replacing rare codons with those preferred by the host’s translational machinery. Traditional codon optimization methods – such as algorithms maximizing the codon adaptation index (CAI) or simply swapping out low-frequency codons – have been successful in many cases and are offered by gene synthesis companies as standard practice^3–5^. However, these conventional approaches have notable limitations: they focus primarily on codon frequency and often neglect other critical sequence features that influence protein expression, such as mRNA secondary structure, codon context, regulatory motifs, and translation kinetics^6–9^. As a result, a sequence optimized purely on codon usage frequencies may still perform suboptimally in the host, and in some instances over-optimization can even reduce protein yield^7,10^. This gap highlights the need for more sophisticated approaches that consider a wider range of factors in gene design.

Orthologous genes – genes in different species that encode the same or similar proteins – provide valuable insights into how nature resolves the challenge of adapting a coding sequence to different organismal contexts^11^. Even between moderately distant species, orthologs typically accumulate numerous non-synonymous substitutions (amino acid-changing mutations) and sometimes small insertions or deletions (indels) over the course of evolution^12^. For instance, when comparing the coding sequences of *Bacillus thuringiensis* genes to their orthologs in *Bacillus subtilis*, only about 57% of codons encode the same amino acid (25% of codons are identical and about 32% are synonymous substitutions where the amino acid is conserved). The remaining 43% of codons differ in amino acid, and in addition, insertions and deletions account for roughly 4% of positions. Similarly high levels of amino acid divergence are observed for more distantly related species (e.g., *Escherichia coli* vs *B. subtilis* orthologs have over 50% non-synonymous codon differences). These observations underscore that adapting a gene to a new species often requires far more extensive rewriting than simple codon swaps. In other words, nature’s solution to cross-species gene transfer is not limited to swapping codons for synonymous alternatives – it also involves adjusting the protein sequence itself when necessary.

These evolutionary insights are consistent with experimental findings: even small differences in a coding sequence can substantially impact expression levels and protein yield in a host cell^13^. For instance, prior studies have shown that synonymous variants of the same gene – differing only by choice of codons – can produce dramatically different amounts of protein in the same host^14,15^. Similarly, using an orthologous gene (a naturally evolved variant from a species adapted to the target host’s clade) sometimes outperforms a naïvely codon-optimized gene in terms of expression. Such observations underscore those factors beyond global codon frequency – such as local sequence context, rare codon placement, and the avoidance of mRNA structural elements or detrimental motifs – can influence translation efficiency and mRNA stability^16^. Therefore, leveraging evolutionary divergence among orthologs offers a promising avenue to inform better codon optimization strategies, by illuminating which sequence changes are well-tolerated or beneficial in a given host background.

At the same time, recent deep learning-based codon optimization methods have emerged to address the shortcomings of purely frequency-based approaches^17,18^. These data-driven models can learn complex relationships in DNA sequences that contribute to expression, going beyond manually defined rules. For example, machine learning models and neural networks have been trained on large collections of synonymous gene variants and expression data to capture the subtle sequence features (e.g. codon pair context, GC content distribution, RNA structural propensity) associated with high protein yields. Deep generative models can design synthetic coding sequences that not only match the host’s codon usage bias but also minimize problematic structures or motifs, effectively learning the language of codons for a given organism. Indeed, several studies have demonstrated improved protein expression using neural network–designed gene sequences compared to those optimized by conventional algorithms^19^. These advances illustrate the power of integrating vast genomic datasets and machine learning to refine codon optimization, enabling what some have called “intelligent” codon optimization that accounts for a multitude of sequence features simultaneously.

Nevertheless, current learning-based codon optimization methods do not explicitly incorporate the evolutionary information contained in orthologous sequences. Evolution has already explored a combinatorial space of coding sequences across diverse organisms; thus, there is an opportunity to harness these natural experiments in combination with deep learning. Here we introduce OrthologTransformer, a novel codon optimization framework that integrates deep learning with evolutionary insights from orthologous gene sequences. OrthologTransformer is built on a Transformer-based architecture, designed to convert a coding sequence from one organism’s genomic context to that of another, effectively emulating how an orthologous gene in the target species might have evolved. By training on known ortholog pairs and large sequence datasets, the model learns to capture host-specific codon preferences, regulatory sequence patterns, and other latent features that contribute to efficient expression in the target organism. In contrast to traditional methods that replace codons in isolation, OrthologTransformer considers the full sequence context and leverages learned evolutionary patterns to propose synonymous substitutions that are context-appropriate and biologically plausible for the host. We hypothesize that this evolution-informed, context-aware approach to codon rewriting will produce genes that achieve higher expression and more consistent reliability in practice.

In this work, we describe the OrthologTransformer model and its application to cross-species gene design. We first validate the model’s performance on converting genes between bacterial species with known orthologs, demonstrating that it can accurately predict the sequence of orthologous genes including their non-synonymous differences. We then showcase a practical case study: the PETase enzyme, which enables bacteria to degrade PET plastic^20^. We used OrthologTransformer to convert the PETase gene from its native bacterium (*Ideonella sakaiensis*) to a target host species (*B. subtilis*), producing a novel gene sequence encoding a putative orthologous enzyme. We synthesized this OrthologTransformer-designed gene and tested it experimentally, finding that the redesigned enzyme retains PET-degradation activity in the *B. subtilis* host. This result provides a proof-of-concept that our model’s predictions are not only computationally plausible but also biologically viable. Overall, OrthologTransformer opens the door to functional gene redesign across species, an advance with broad implications for synthetic biology, biotechnology, and our understanding of evolutionary protein adaptation.

## Results

### A deep learning model for orthologous gene conversion

To enable gene sequence adaptation beyond synonymous changes, we developed OrthologTransformer, a sequence-to-sequence model that converts a coding DNA sequence from a source species into a sequence that could be its ortholog in a target species. We trained the model on a large collection of orthologous gene pairs drawn from diverse bacteria. Each training example consisted of a coding sequence from species A as input and the corresponding coding sequence from species B as the target output. In training the model to map one sequence to the other, we implicitly teach it the DNA-level conversions that connect orthologs across those two species.

Our model uses the Transformer architecture, as illustrated in Figure 1a, which is well-suited for modeling long sequences with complex dependencies^21,22^. We represented gene sequences at the level of codons (3-base tokens) rather than individual nucleotides. This codon-level representation drastically reduces sequence length and aligns with the biological notion of codon substitution. It also allows the model to capture synonymous vs non-synonymous changes explicitly – a single token change can alter an amino acid or not, depending on whether the codon is replaced by a synonym or a different amino-acid codon. We included special tokens to denote start and stop codons, and the model is capable of handling insertions or deletions by producing an output sequence that is longer or shorter than the input. In essence, the Transformer learns to “edit” the input codon sequence, inserting or removing codon tokens and changing codons as needed to produce the output sequence. Because the training ortholog pairs are naturally aligned in terms of protein function, the model learns to make changes that preserve function (as orthologs do) rather than random destructive mutations.

**Figure 1.**
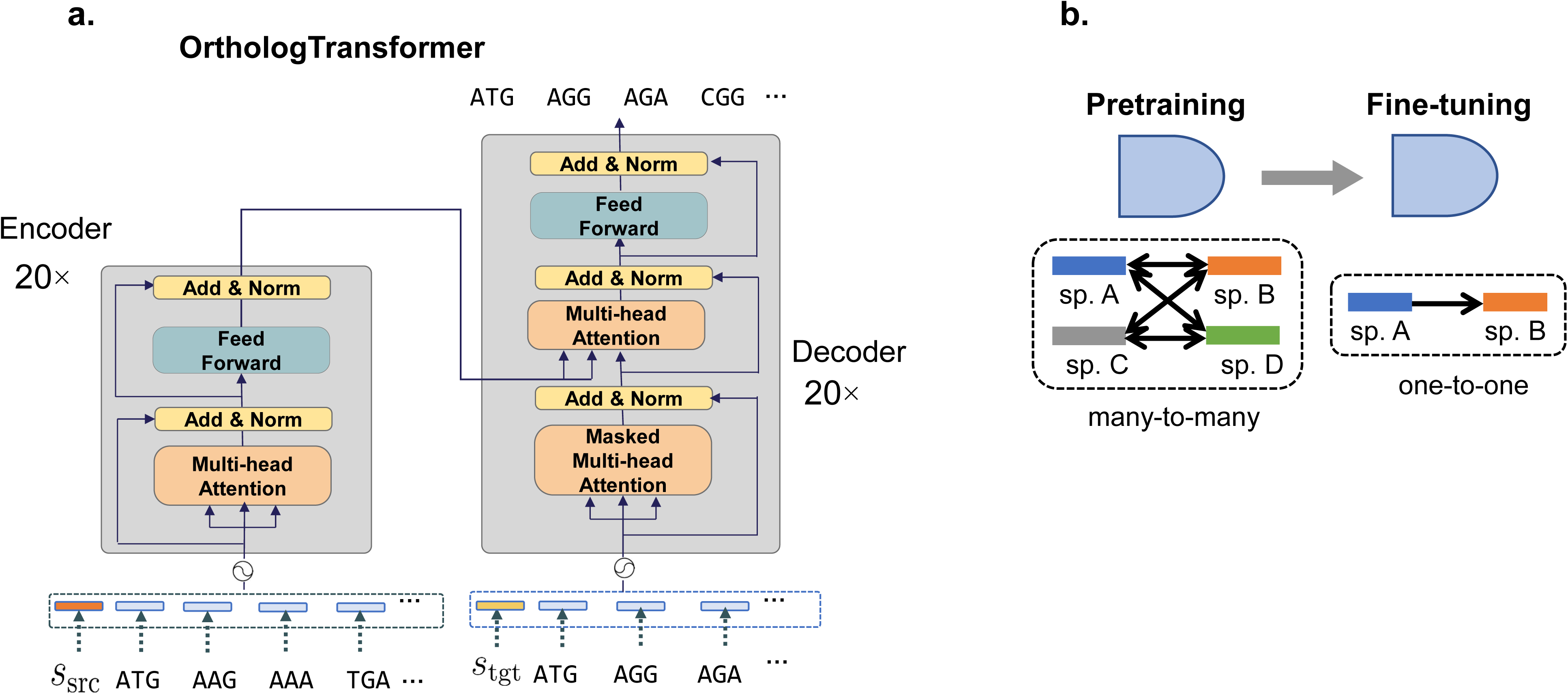
Architecture of OrthologTransformer. **a.** The model features a 20-layer encoder-decoder structure, with each layer equipped with Add & Normalization layers and Multi-head Attention mechanisms. Species signals (*s*_*src*_, *s*_*tgt*_) are prepended to the input sequence, enabling species-specific sequence conversion. **b.** OrthologTransformer employs a two-stage learning approach consisting of pretraining and fine-tuning. In the pretraining phase, the model learns general sequence conversion patterns from many-to-many orthologous relationships across multiple species. In the fine-tuning phase, the model is specialized for specific one-to-one species pair conversions using targeted training data.

During training, we conditioned the model on the specification of the target species. In practice, this was achieved by prepending a special token indicating the target species to the input sequence. This way, a single model can handle conversion to many possible target species (Figure 1b), learning each species’ particular codon usage and typical amino acid adaptations. The final trained OrthologTransformer can take an input gene and a specified target species and generate a predicted orthologous gene sequence for that species. We note that the model is not simply doing a multi-species codon optimization; it has free reign to alter amino acids or gene length if the training data suggest that such changes commonly occur for that gene or between those species.

### Validation on known orthologous genes

To evaluate OrthologTransformer, we first tested how well it could predict actual ortholog sequences between species in our dataset. For a given pair of species (A and B), we withheld some known ortholog pairs from training, and then asked the model to convert those genes from A to B. We then compared the model’s output to the true orthologous sequence in species B. We trained OrthologTransformer on a large corpus of orthologous gene pairs to enable cross-species DNA converton. The model’s conversion accuracy (the fidelity of converting a gene from source to target species coding) improved as the training dataset expanded. As shown in Table 1, our training datasets ranged from one-to-one ortholog pairs between 2 species (658 sequence pairs) to many-to-many ortholog relationships across 2,138 species (over 5.6 million sequence pairs from the OMA database^23^). This progressive expansion of training data yielded remarkable improvements in ortholog conversion accuracy, as demonstrated in Figure 2, where the sequence identity between generated sequence and target sequence dramatically increased from 0.15 when using the smallest dataset (2 species) to 0.40 when leveraging the full 2,138-species dataset. By exposure to data from more diverse bacterial species, the model learned universal patterns in inter-species gene adaptation. Additionally, we found that for small-scale datasets, alignment processing of training data and model fine-tuning were effective for improving accuracy, while for large-scale datasets, the natural diversity of sequences itself provided sufficient learning signals, achieving high accuracy even without alignment processing.

**Figure 2.**
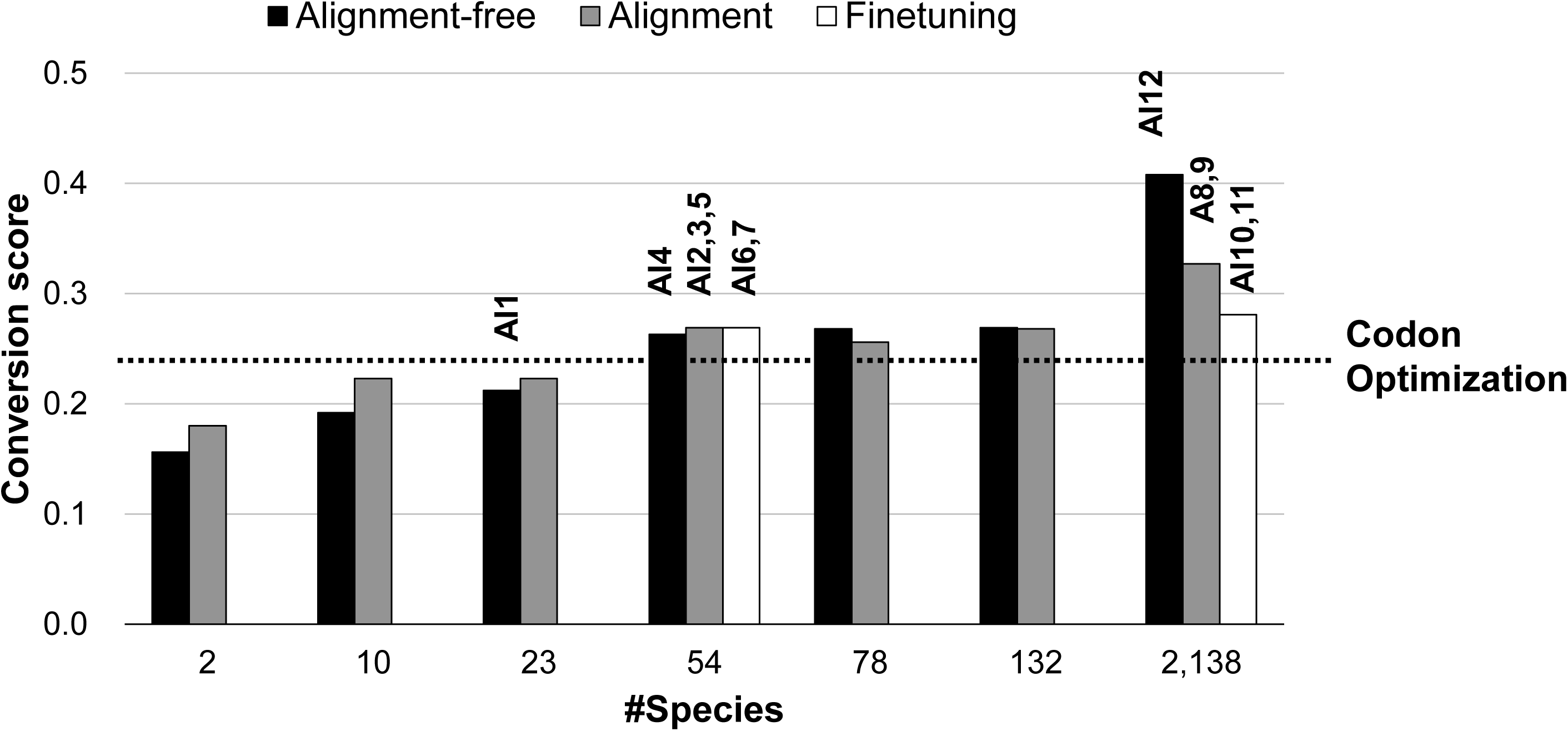
Performance improvement with increasing training dataset scale. Evaluation of accuracy when converting from *I. sakaiensis* to *B. subtilis*. Each bar graph shows the conversion score (sequence identity) between the generated sequence and the target sequence at different dataset scales (ranging from 2 to 2,138 bacterial species). Experiments were conducted using three approaches: models without alignment processing (Alignment-free, black), models with alignment processing (Alignment, gray), and models with fine-tuning (Finetuning, white). The numbers within the graph indicate the actual number of sequence pairs used in each dataset. The vertical axis represents the sequence identity between the generated sequence and the target sequence (range 0 to 1), while the horizontal axis shows the number of bacterial species used.

**Table 1.** Training datasets for OrthologTransformer.

| Type | #species | #sequence pairs | #DNA sequences | Database |
| --- | --- | --- | --- | --- |
| one-to-one | 2 | 658 | 1,014 | RefSeq+Orthofinder<sup>45</sup> |
| many-to-many | 10 | 12,882 | 40,223 |  |
| many-to-many | 23 | 67,881 | 120,119 |  |
| many-to-many | 54 | 129,297 | 283,438 |  |
| many-to-many | 78 | 201,330 | 633,309 |  |
| many-to-many | 132 | 252,449 | 701,232 |  |
| many-to-many | 2,178 | 4,971,060 | 4,411,521 | OMA<sup>23</sup> |

Training datasets used for OrthologTransformer development. The table shows the type of orthologous relationships (one-to-one or many-to-many), number of species used, number of sequence pairs, total DNA sequences, and the database source for each training set configuration.

To further illustrate OrthologTransformer’s capabilities, we examined both codon-level and amino acid-level adaptations across various bacterial species. As Table 2 shows, the sequence identity between the generated sequence and the target sequence was significantly improved compared to the original source sequence, particularly between species exhibiting vastly different genomic or physiological characteristics. For example, the identity to the target sequence doubled from 0.221 (original source sequence) to 0.408 (generated sequence) when converting between *B. subtilis* (43.5% GC content) and *I. sakaiensis* (66.7% GC content). Even more notably, between *L. lactis* and *T. thermophilus*, which differ substantially in optimal growth temperatures (30°C versus 65°C), the sequence identity increased more than threefold – from 0.157 (original source sequence) to 0.472 (generated sequence). Across all tested species pairs, the generated sequences consistently displayed higher similarity to the target sequences than did the original source sequences, clearly illustrating the model’s ability to effectively enhance sequence adaptation for diverse species pairs.

**Table 2.** Performance comparison of OrthologTransformer across bacterial species pairs.

| Species pair | Codon sequence identity |  |  |  | Amino acid sequence similarity |  |
| --- | --- | --- | --- | --- | --- | --- |
|  | generated vs target |  |  | source vs target | generated vs target | source vs target |
|  | Ortholog Transformer | Codon optimization | CodonTransformer<sup>24</sup> |  |  |  |
| *I.sakaiensis*→*B. subtilis* | 0.408 | 0.265 | 0.286 | 0.221 | 2.862 | 1.693 |
| *Streptomyces*→*B. subtilis* | 0.376 | 0.265 | 0.290 | 0.218 | 2.581 | 1.762 |
| *Synechococcus*→*E.coli* | 0.740 | 0.321 | 0.328 | 0.279 | 4.070 | 1.844 |
| *P.putida*→*E.coli* | 0.690 | 0.340 | 0.344 | 0.323 | 3.407 | 2.413 |
| *M.tuberculosis*→*E.coli* | 0.697 | 0.300 | 0.308 | 0.266 | 3.577 | 1.623 |
| *T.thermophilus*→*L. lactis* | 0.472 | 0.326 |  | 0.157 | 2.673 | 1.760 |
| *P.fluorescens*→*R.leguminosarum* | 0.464 | 0.384 |  | 0.331 | 2.093 | 1.987 |
| *D.radiodurans*→*S.typhimurium* | 0.633 | 0.315 |  | 0.285 | 3.042 | 1.720 |
| *L.acidophilus*→*A.vinelandii* | 0.429 | 0.326 |  | 0.193 | 2.520 | 1.495 |

This table shows codon sequence identity and amino acid sequence similarity between the source sequence and sequences generated by OrthologTransformer, frequency-based codon optimization, and CodonTransformer<sup>24</sup>, each compared against the corresponding target sequence. For CodonTransformer, comparisons are shown only for species pairs included in the publicly available trained model. Regarding amino acid sequence similarity, comparisons were performed only for sequences generated by OrthologTransformer, since frequency-based codon optimization and CodonTransformer involve only synonymous codon substitutions, and thus no amino acid substitutions occur. Across all evaluated pairs, OrthologTransformer consistently achieves higher codon identity than both frequency-based codon optimization and CodonTransformer, while preserving high amino acid similarity.

Turning to comparisons with existing methods, OrthologTransformer achieves approximately a 1.7-fold improvement in average codon-sequence identity compared to conventional codon optimization across all species pairs. This suggests that synonymous-only optimization is insufficient to meet the contextual and evolutionary demands of host adaptation. CodonTransformer, a recent deep learning model, uses a Transformer architecture to optimize synonymous codon choices in a context-aware manner, and is trained on large multispecies datasets^24^. However, in the subset of five species pairs where CodonTransformer was evaluated, OrthologTransformer still maintains a clear advantage, delivering an average improvement of approximately 1.8-fold. These findings highlight that by incorporating non-synonymous substitutions and indels, OrthologTransformer enables a more advanced and effective form of gene adaptation than existing synonym-focused methods.

Beyond mere sequence identity, OrthologTransformer preserves functional meaning through appropriate amino acid adaptations. Table 2’s BLOSUM score analysis reveals significant functional conservation improvements at the amino acid level between various species pairs. Notably, these substitutions often involve changes to amino acids with similar chemical properties (conservative substitutions), suggesting the model learned to prefer functionally mild changes when altering protein sequences. For thermophilic adaptations, the model appropriately adjusted proportions of arginine and lysine, which are critical for thermal stability. These BLOSUM score improvements indicate that our model has not simply optimized codon usage but has learned to capture the underlying biological principles governing protein adaptation across different cellular environments. This functional preservation is crucial for maintaining enzymatic activity while adapting genes to new host organisms. Detailed analysis of individual ortholog pairs across diverse bacterial species revealed that OrthologTransformer successfully improved sequence adaptation in most cases, as demonstrated by the comparison of pre-conversion versus post-conversion accuracy scores (Supplementary Figures 1 and 2). Another noteworthy aspect is that OrthologTransformer can achieve ortholog conversions that comprehensively capture the genetic characteristics specific to target bacterial species, not just increase sequence identity. The analysis in Figure 3 revealed that OrthologTransformer can simultaneously optimize important sequence characteristics such as GC content and CAI (Codon Adaptation Index). Through learning on large-scale datasets, the model acquired universal patterns of gene adaptation, and through fine-tuning to specific pairs, it could further enhance species-specific adaptations.

**Figure 3.**
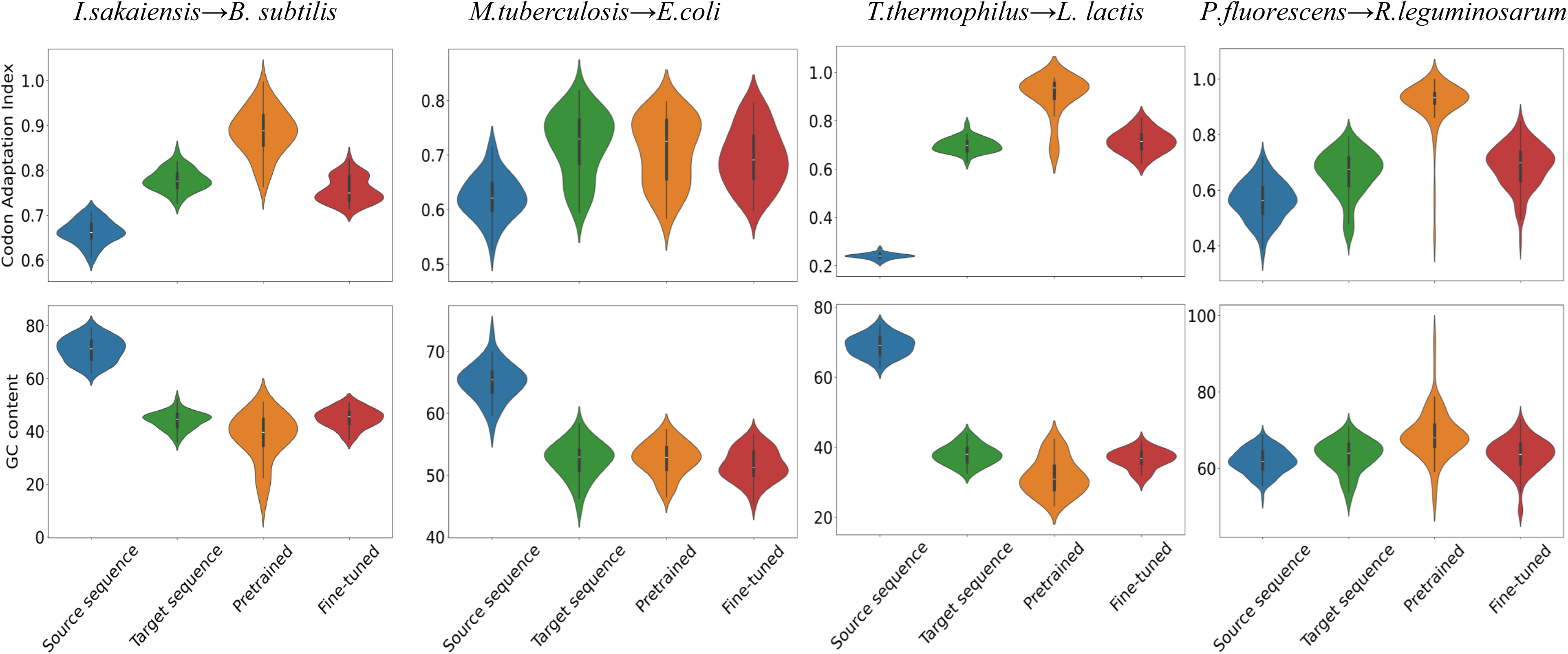
Comparison of CAI and GC content distribution across different bacterial species pairs. Each panel shows ortholog conversion results for different bacterial species pairs. The upper portion of each panel displays violin plots representing the distribution of the Codon Adaptation Index (CAI), while the lower portion shows the distribution of GC content. Blue represents the source sequences, green indicates the target sequences, orange corresponds to sequences generated by the pretrained model, and red denotes sequences generated by the fine-tuned model.

### Designing a PETase ortholog for B. subtilis

While validation against known orthologous genes is useful, the ultimate goal of OrthologTransformer is to enable *de novo* gene adaptation for cases where the target species lacks a known ortholog. To demonstrate this capability, we applied OrthologTransformer to the PETase enzyme gene from *Ideonella sakaiensis*. PETase is a PET (polyethylene terephthalate) plastic-degrading enzyme originally identified in *I. sakaiensis*, a bacterium that can break down PET as a carbon source^20,25^. There is great interest in expressing PETase in other host microbes (such as *E. coli* or *B. subtilis*) to harness or improve plastic degradation, but the native PETase gene might not express optimally outside *I. sakaiensis*. Moreover, *I. sakaiensis* is a relatively distant organism; common lab hosts like *B. subtilis* do not have a direct ortholog of PETase. This makes PETase an excellent test case for OrthologTransformer’s ability to generate a functional orthologous sequence from scratch.

### In silico design

We input the PETase coding sequence (from *I. sakaiensis*) into OrthologTransformer, specifying *B. subtilis* as the target species. OrthologTransformer produced an *B. subtilis*-adapted version of the PETase gene. We attempted to convert the PETase gene using various design approaches (Table 3). We systematically varied experimental conditions, including the scale of the training dataset (ranging from 23 to 2,138 species), the presence or absence of alignment processing, the introduction of Monte Carlo Tree Search (MCTS), and fine-tuning. Our MCTS strategy carries out multi-objective sequence optimization that simultaneously enforces low GC content – to facilitate DNA synthesis – and stable mRNA secondary structures by minimising folding free energy, thereby supporting higher translational efficiency. Consequently, we designed and evaluated a total of 12 sequences, termed “AI-designed sequences" (denoted AI1 – AI12).

**Table 3.**
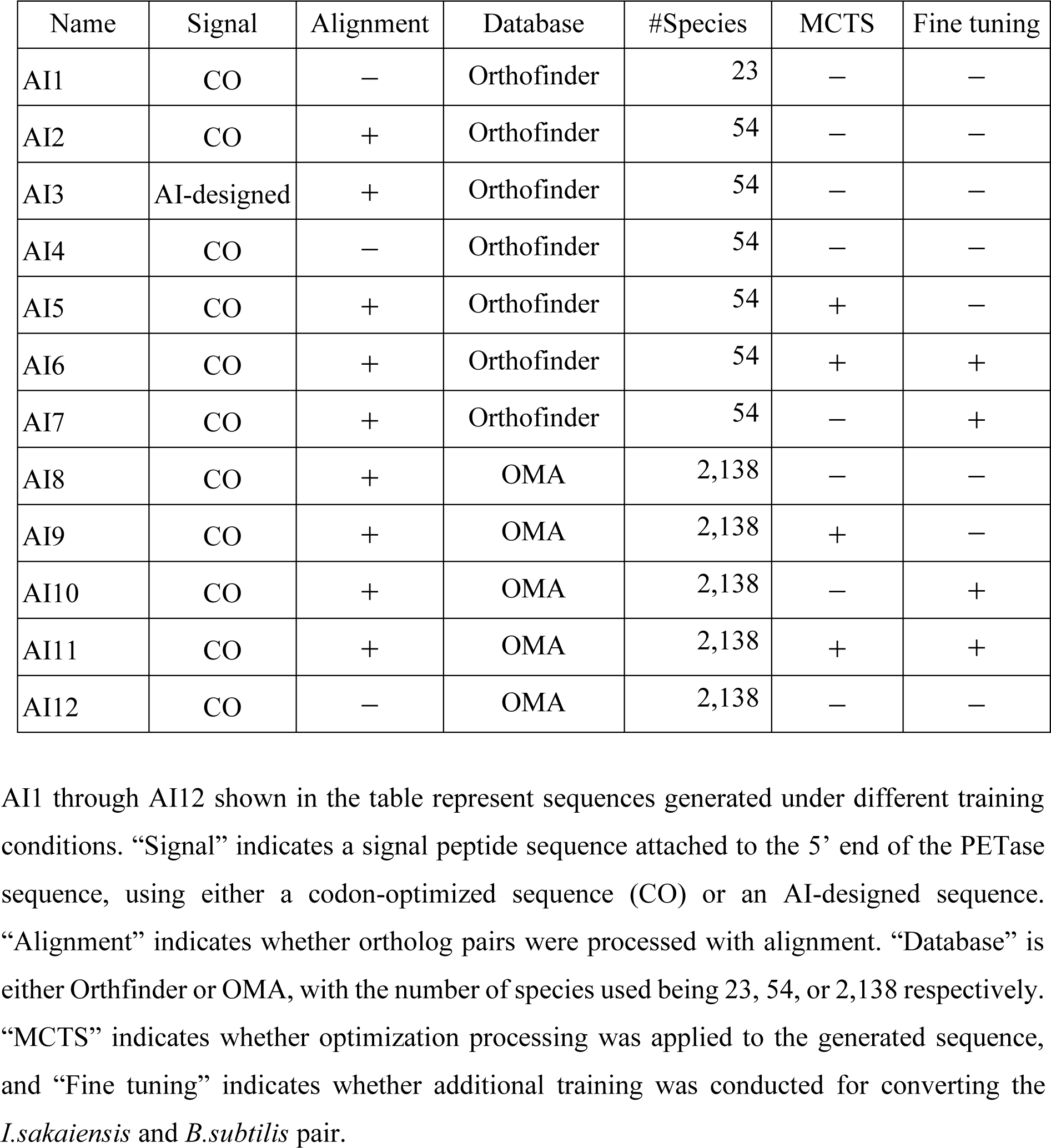
Experimental conditions for PETase sequence design variants.

| Name | Signal | Alignment | Database | #Species | MCTS | Fine tuning |
| --- | --- | --- | --- | --- | --- | --- |
| AI1 | CO | — | Orthofinder | 23 | — | — |
| AI2 | CO | + | Orthofinder | 54 | — | — |
| AI3 | AI-designed | + | Orthofinder | 54 | — | — |
| AI4 | CO | — | Orthofinder | 54 | — | — |
| AI5 | CO | + | Orthofinder | 54 | + | — |
| AI6 | CO | + | Orthofinder | 54 | + | + |
| AI7 | CO | + | Orthofinder | 54 | — | + |
| AI8 | CO | + | OMA | 2,138 | — | — |
| AI9 | CO | + | OMA | 2,138 | + | — |
| AI10 | CO | + | OMA | 2,138 | — | + |
| AI11 | CO | + | OMA | 2,138 | + | + |
| AI12 | CO | — | OMA | 2,138 | — | — |

AI1 through AI12 shown in the table represent sequences generated under different training conditions. “Signal” indicates a signal peptide sequence attached to the 5’ end of the PETase sequence, using either a codon-optimized sequence (CO) or an AI-designed sequence. “Alignment” indicates whether ortholog pairs were processed with alignment. “Database” is either Orthofinder or OMA, with the number of species used being 23, 54, or 2,138 respectively. “MCTS” indicates whether optimization processing was applied to the generated sequence, and “Fine tuning” indicates whether additional training was conducted for converting the *I.sakaiensis* and *B.subtilis* pair.

Importantly, OrthologTransformer not only substituted codons but also introduced changes at the amino acid level suitable for the host environment. As illustrated in Figure 4, these modifications involved varying degrees of insertions, deletions, synonymous substitutions, and non-synonymous substitutions across the 12 AI-designed sequences. The extent of these changes varied dramatically among the different versions: from minimal modifications in AI8 (no changes) to extensive remodeling in AI4 (160 insertions, 139 deletions, 72 synonymous substitutions, and 30 non-synonymous substitutions). Despite such extensive sequence modifications, structural predictions showed that key functional domains remained intact.

**Figure 4.**
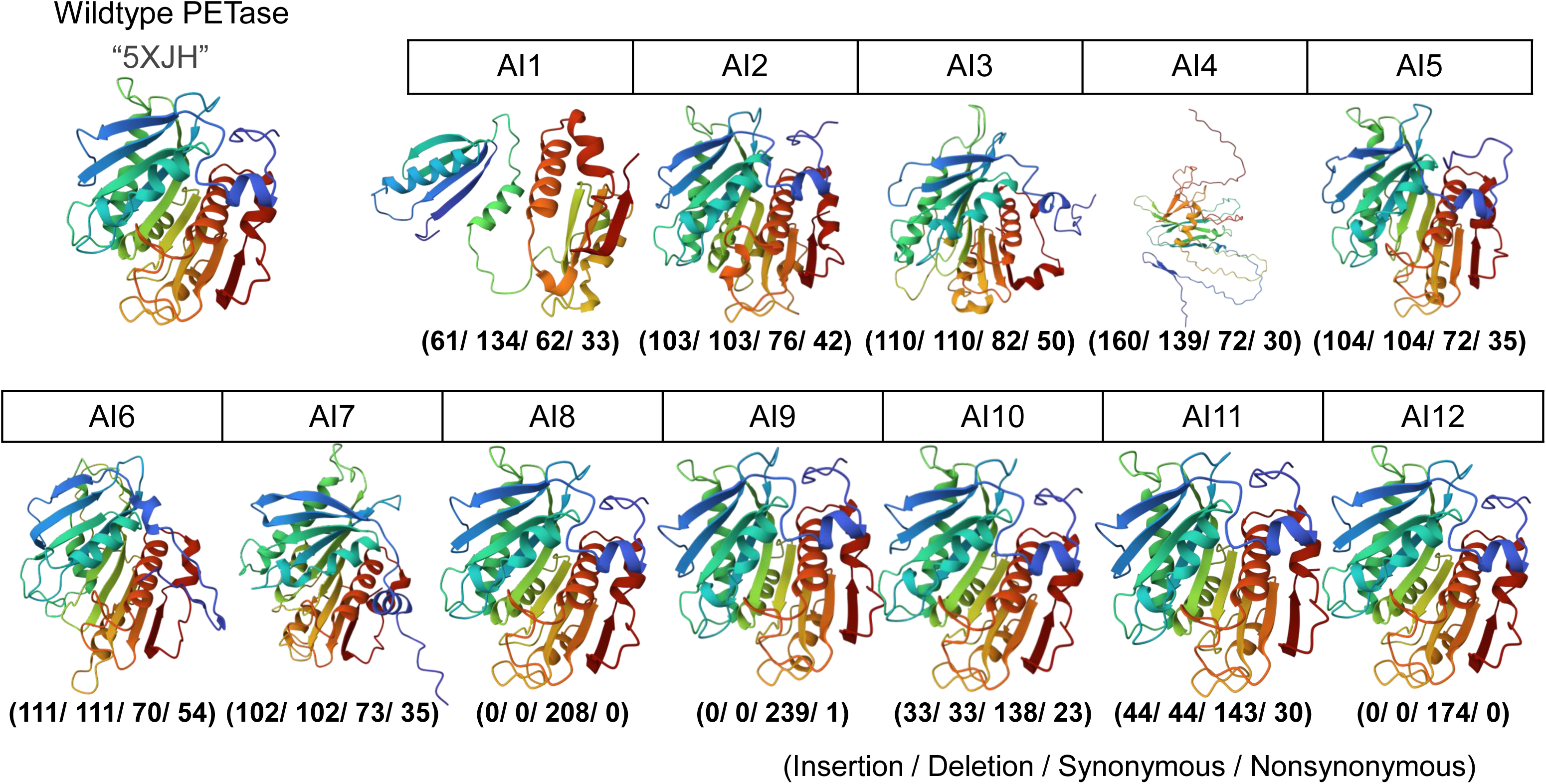
Structural prediction and sequence modifications of AI-designed PETase variants. OrthologTransformer generated twelve different PETase variants (AI1–AI12) with various degrees of sequence modifications. The wild-type PETase structure (PDB entry 5XJH) is shown on the left for reference. The numbers below each structure indicate the modifications introduced in each variant: (insertions/deletions/synonymous substitutions/nonsynonymous substitutions).

The AI9 sequence demonstrated particularly groundbreaking results by combining training on a large-scale dataset with multi-objective optimization via MCTS. MCTS implemented a strategic two-stage optimization: first controlling GC content before transitioning to RNA secondary structure stabilization. This approach successfully resolved the typical trade-off between reducing GC content and improving RNA structure stability. As illustrated in Figure 5, AI9 achieved an optimal balance with 37.0% GC content, -281 kcal/mol RNA secondary structure stability, and a 0.98 TM score. These metrics represent substantial improvements over both wild-type and conventional codon-optimized sequences, fulfilling the three critical sequence design requirements: synthesis feasibility, mRNA stability, and protein structure conservation. Thus, the newly designed PETase sequence can be viewed as an “AI-suggested ortholog” of the original PETase gene, tailored for *B. subtilis*.

**Figure 5.**
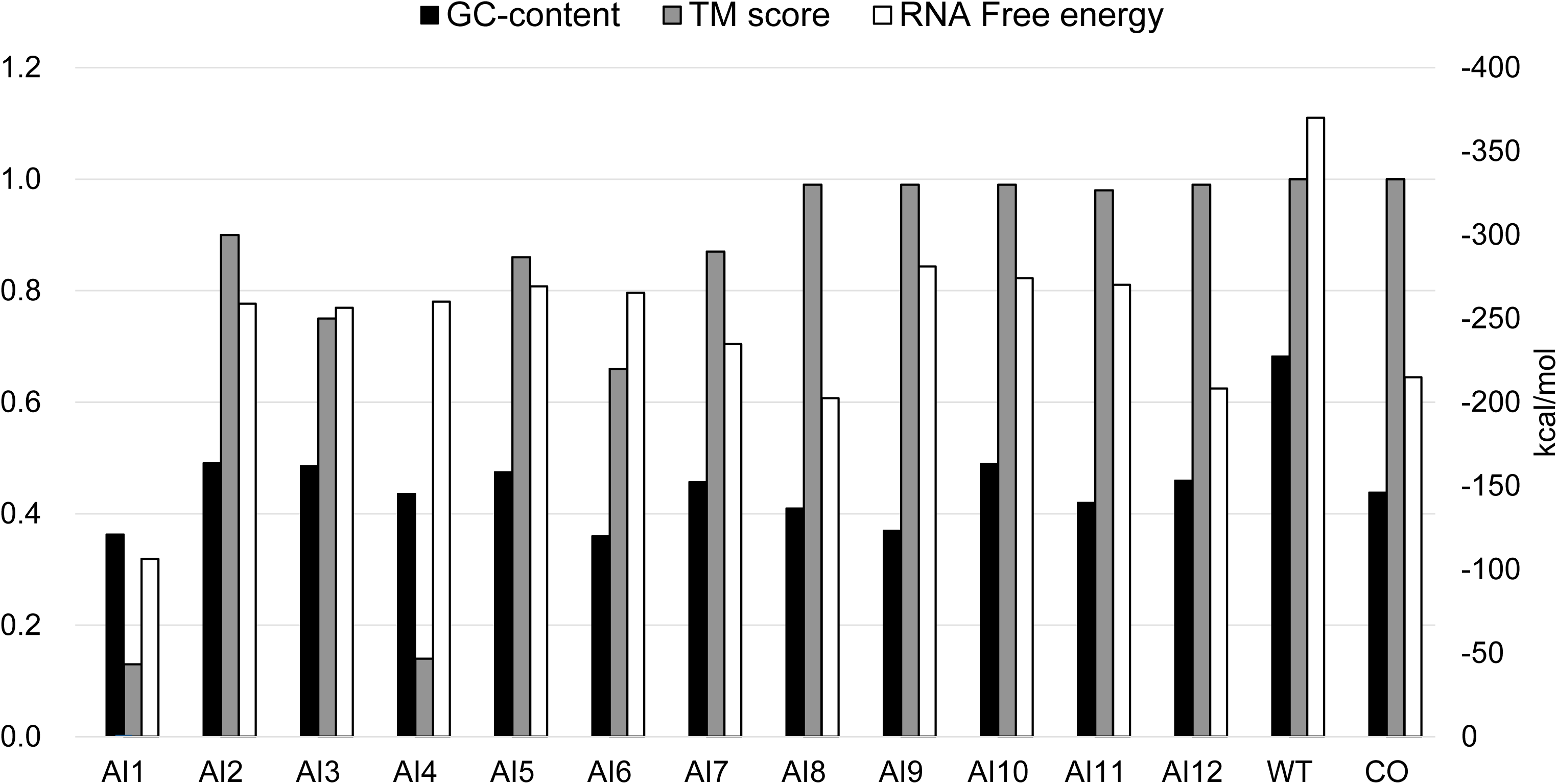
Sequence properties of PETase variants. Comparison of GC content, TM score, and RNA secondary structure free energy among AI-designed PETase variants (AI1–AI12), wild-type (WT), and codon-optimized (CO). AI-designed variants, particularly those trained on larger datasets, achieved superior balance across all three parameters, demonstrating the effectiveness of multi-objective optimization over conventional codon optimization.

### Experimental expression and PET degradation activity assay

We synthesized the AI-designed (OrthologTransformer-designed) PETase genes (AI1 – AI12) and cloned them into an expression vector for *B. subtilis*. We included two controls: the wild-type *I.sakaiensis* PETase gene (denoted WT) and a codon-optimized PETase gene for *B. subtilis* (denoted CO), which had the same amino acid sequence as wild-type PETase but with every codon replaced by the most preferred *B. subtilis* synonymous codon for that amino acid. All constructs were engineered with an N-terminal secretion signal peptide for Sec-pathway export and a C-terminal 6×His tag for detection (Supplementary Figure 5).

Initially, we assessed the transcription and translation of the AI-designed genes and two controls in *B. subtilis*.

#### mRNA full-length transcription

We first confirmed that each strain transcribed the full-length PETase ORF. PCR using primers spanning the 5’ and 3’ ends of the PETase coding sequence (including the signal peptide region and His-tag) yielded the expected product in all engineered strains (Figure 6), demonstrating successful transcription across the entire coding region, including both the 5’ signal peptide and 3’ His-tag sequences. Thus, the AI-designed sequences were genomically stable and correctly processed by the host transcription machinery.

**Figure 6.**
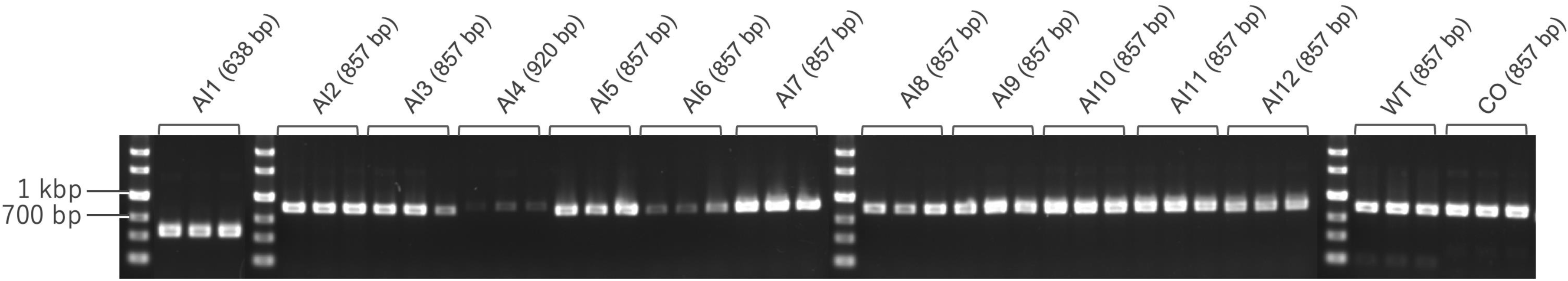
Confirmation of full-length transcription of PETase gene. Full-length transcription of the PETase gene was confirmed in all recombinant strains (n=3). Primers were designed for the 5’ and 3’ ends of the PETase gene, including the signal and His tag sequences, which were common across all recombinant strains.

#### Transcription levels (qPCR)

We quantified PETase mRNA levels by quantitative real-time PCR (qPCR), normalizing to the *B. subtilis* housekeeping gene *gyrA*. All AI-designed strains produced substantial PETase transcripts, though with variation across designs. Several AI-designed strains, such as those harboring AI1, AI2, and AI8, exhibited high transcript abundance, either comparable to or exceeding that of the codon-optimized control. Notably, the strain harboring AI2 showed among the highest PETase mRNA levels, statistically indistinguishable from those of the two controls (WT and CO) under inducing conditions (Figure 7). These differences might reflect how certain synonymous choices or nucleotide changes influenced mRNA stability or transcription efficiency^10^.

**Figure 7.**
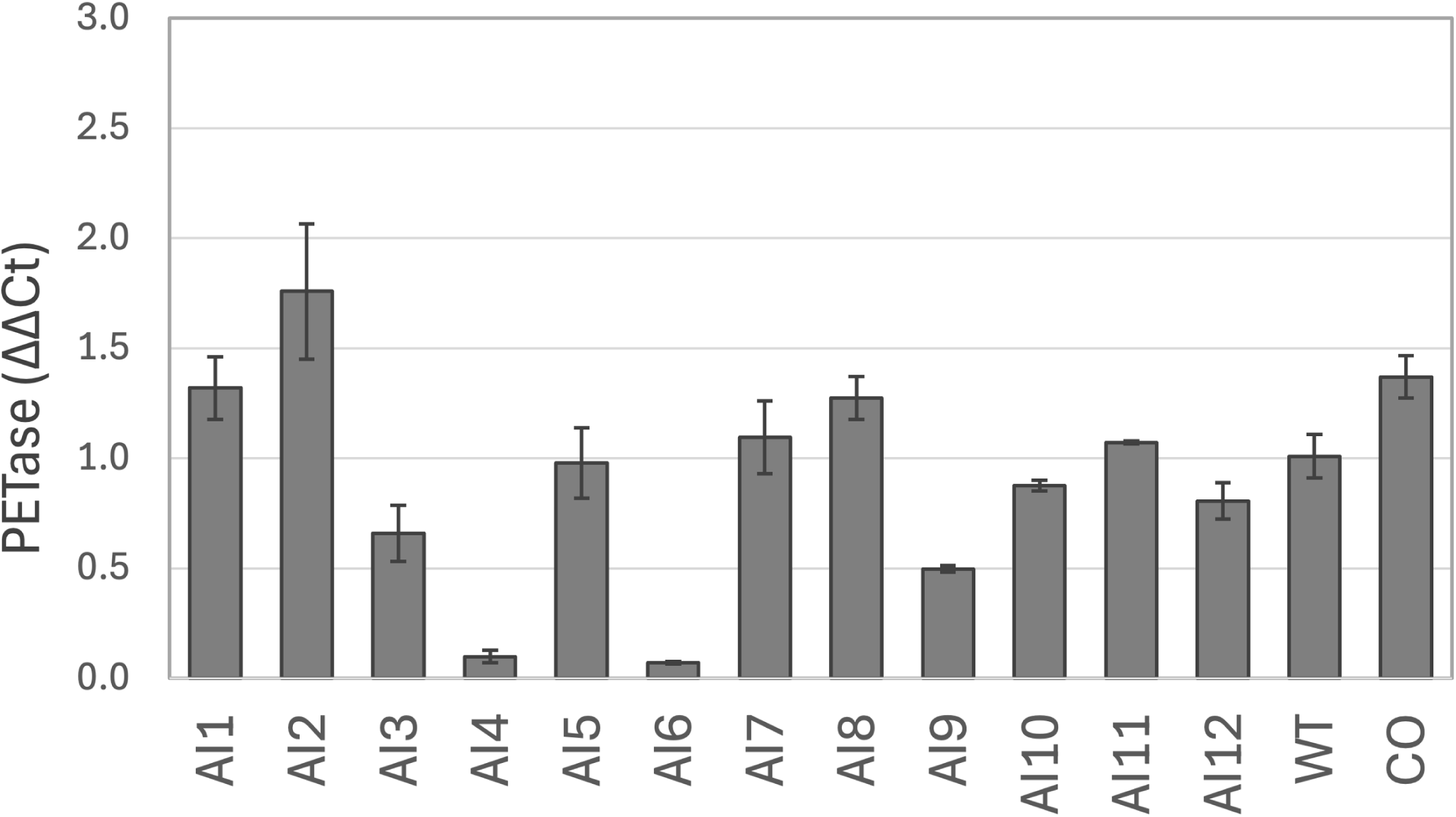
Quantification of PETase gene transcription by qPCR. The transcription levels of the PETase gene were quantified relative to the housekeeping gene *gyrA* of *B. subtilis* using qPCR (n=3).

#### Protein expression and secretion

We evaluated PETase protein production by western blot analysis. Culture supernatants and cell pellets were collected from *B. subtilis* cultures following induction, and PETase was detected using an anti-His-tag antibody. The detection of PETase was performed using the BCIP/NBT method, which revealed distinct bands corresponding to PETase (∼30 kDa) in the supernatant fractions of multiple engineered strains (Figure 8). Notably, the AI-designed variants AI8, AI9, and AI12 demonstrated robust PETase secretion. The presence of PETase in the media confirms that the signal peptide functioned and the enzyme was exported out of the cells (a crucial feature for PET degradation, since PET is extracellularly located as a solid substrate). No immunoreactive bands were observed in the empty-vector negative control. Among the variants tested, AI8 and AI9 showed clear PETase signals on the blot and demonstrated exceptional effectiveness within the host’s expression system.

**Figure 8.**
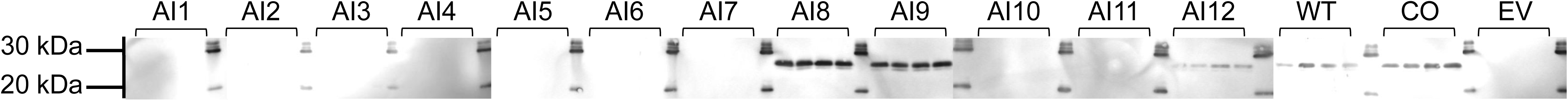
Western Blotting for PETase detection by BCIP/NBT. Culture supernatants were collected after 12 hours cultivation, and western blotting was conducted by 6xHis-tag antibody (n=4). While the bands of WT and AI12 were faint, sharp bands were detected at AI8 and AI9.

#### PET degradation activity of AI-designed PETase

Finally, we tested the functional activity of each enzyme using a PET degradation assay. In this assay, *B. subtilis* cells expressing PETase were incubated with a film of additive-free PET plastic, and the breakdown products (terephthalic acid (TPA), mono(2-hydroxyethyl) terephthalate (MHET) and Bis(2-hydroxyethyl) Terephthalate (BHET)) were measured over time by HPLC method (Supplementary Figure 4). The results confirmed that the AI-designed PETase is functional. *B. subtilis* cells harboring the AI1 – AI12 genes exhibited clear PET degradation activity, comparable to cells with the codon-optimized PETase CO. We evaluated PET film degradation by the engineered strains in liquid culture over 7 days. PET hydrolysis was monitored by measuring its soluble breakdown products (TPA, MHET, and BHET) in the culture supernatant on days 1, 2, 3, and 7. PET hydrolysis was monitored by measuring its soluble breakdown products (TPA, MHET, and BHET) in the culture supernatant on days 1, 2, 3, and 7. Due to PETase’s known endo-activity, which predominantly generates MHET, the accumulation of MHET serves as the expected indicator of PETase activity in this assay system. MHET was detected as early as the second day in some engineered strains (AI3, AI4, AI9, AI10, AI12, and WT), and by the third day MHET had accumulated in all PETase-expressing cultures (Figure 9a). In contrast, no TPA or BHET was detected in any of the cultures throughout the 7-day incubation (Supplementary Figure 6), indicating that MHET was the primary hydrolysis product. Notably, the AI9 variant stood out by producing roughly three-fold more MHET than any other strain by day 3, reflecting significantly higher PET-degrading activity (p < 0.05; Figure 9b). This result indicates that AI9 has the strongest PET hydrolytic activity among the variants tested.

**Figure 9.**
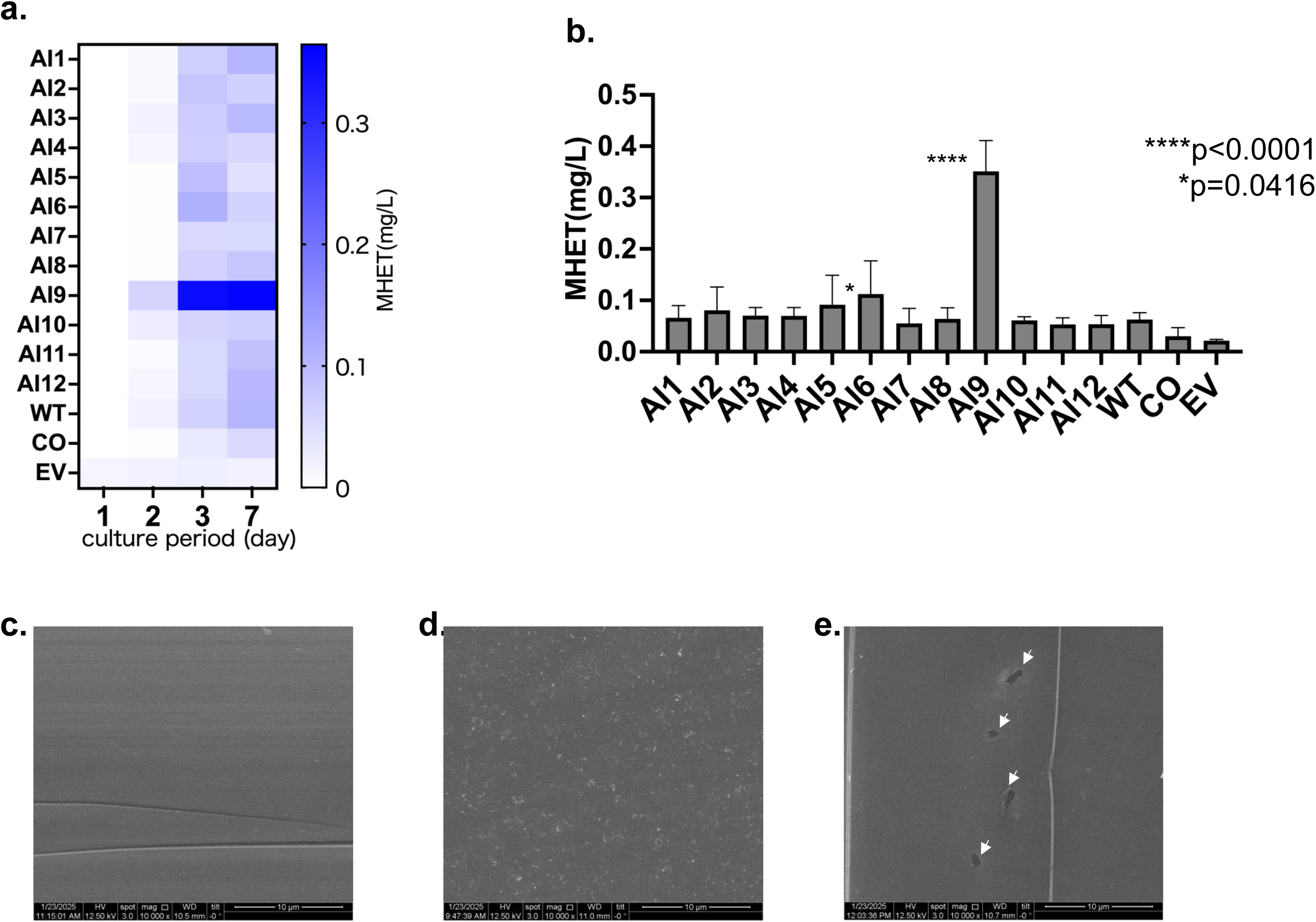
Evaluation experiment using PET film: Quantification of PET degradation products by HPLC and scanning electron microscopy (SEM) of the PET films. **a.** MHET hydrolyzed by PETase from PET film was measured during cultivation of engineered strains over 7 days. **b.** MHET hydrolyzed by PETase from PET film was measured during cultivation of engineered strains on 3rd day. All experiments were performed at least in triplicate. Error bars indicate standard deviation. *p < 0.05, ****p < 0.0001 (Ordinary one-way ANOVA). **c-d**. Degradation of PET film treated with (c) EV, (d) AI9, (e) AI6, as visualized by SEM. Arrow shows the changes.

Consistent with the MHET measurements, scanning electron microscopy (SEM) of the PET films after 7 days revealed the most extensive surface erosion in the AI9 treatment. The PET film incubated with the negative control (PET incubated with the empty-vector strain, denoted EV) remained mostly intact, showing a smooth surface with no obvious damage (Figure 9c). In contrast, the film recovered from the AI9 culture displayed large cavities and “holes” approximately the size of bacterial cells (1-2 µm) penetrating into the film (Figure 9d), suggesting that colonies of *B. subtilis* on the film had extensively degraded the plastic directly beneath them. large, cell-sized pits perforating the surface (Figure 9d). The AI6 strain showed only slight surface pitting (small irregular depressions) (Figure 9e), consistent with its lower MHET yield. The other PETase strains also caused some surface roughening and etched patterns (Supplementary Figure 7), whereas the control remained smooth. These qualitative observations align with the quantitative data, reinforcing that the AI9 strain achieves the most pronounced PET degradation among the strains examined.

In summary, the AI-designed PETase variants were functional and capable of degrading PET *in vivo*. Among them, AI9 was the top performer, producing the most MHET and the most pronounced PET film decomposition. The improved activity of AI9 correlates with its optimized sequence features (balanced GC, stable mRNA) that likely drove higher protein production. These results provide a compelling proof-of-concept that AI-guided gene design can directly convert into enhanced biochemical function in a target host.

## Discussion

OrthologTransformer represents a significant evolution of the concept of gene optimization for heterologous expression. By moving beyond purely synonymous alterations, it addresses a major gap left by traditional codon optimization methods. Standard codon optimization is inherently limited: it “optimizes” a gene only in terms of codon usage bias, while freezing the protein sequence^1,4^. This strategy cannot account for situations where the protein sequence itself may benefit from change – for example, to better fold or function in the biochemical milieu of the host organism^26^. OrthologTransformer tackles this challenge by leveraging the information encoded in natural orthologs.

Our results highlight several key advantages of this approach. First, OrthologTransformer can capture codon context effects that go beyond individual codon usage frequencies. Because it considers the full sequence (and even flanking sequences, if included) when generating output, it can, for instance, avoid creating problematic mRNA secondary structures or inadvertent motifs (such as premature poly-A signals or rare restriction sites) that simpler codon optimizers might miss. Second, and most importantly, OrthologTransformer introduces non-synonymous substitutions and indels in a controlled, learned manner. These changes are not random; they are grounded in patterns seen between orthologous genes. For example, if a certain alanine in a protein tends to be replaced by valine in a related species’ orthologs, the model may do the same when converting a new gene. Such changes could impart subtle improvements in stability or interactions with species-specific partner proteins or chaperones. In the *Bacillus* and *E. coli* comparisons, we saw that a large fraction of amino acid differences between species could be correctly predicted by the model, which is a strong indication that the model is learning general rules of protein adaptation.

Our results demonstrate that OrthologTransformer can bridge the genomic and functional gap between species, enabling a gene from one organism to be re-coded for another organism’s cellular machinery while retaining its function. In this study, a PETase gene from *I. sakaiensis* (a Gram-negative bacterium) was successfully converted into a form that *B. subtilis* (a Gram-positive bacterium) can express at high levels. Achieving this required more than simple codon optimization; by leveraging deep learning trained on thousands of evolutionary variants, OrthologTransformer captured subtle sequence features that make a gene “feel native” to the host genome. The dramatic improvement in PETase expression and activity in *B. subtilis* – especially by the AI9 sequence – highlights the power of this approach.

The PETase case also underscores the value of preserving protein structure while altering sequence. We intentionally did not introduce amino acid substitutions in the final designs (aside from any that the model may have implicitly suggested and we validated to not harm activity). By maintaining the enzyme’s amino acid sequence (and thus active site geometry), we ensured that any activity differences were due to expression level rather than intrinsic catalytic changes. Even so, we note that nature’s orthologs sometimes have amino acid differences that confer advantages (e.g. improved stability). In principle, OrthologTransformer can be expanded – given sufficient training data on functional homologs – to actively propose beneficial amino-acid substitutions, thereby unifying protein engineering with gene optimization. Indeed, introducing only a handful of point mutations has been shown to markedly enhance the thermostability and hydrolytic activity of PETase^27,28^, suggesting that layering such residue-level edits onto our host-adapted designs could unlock even greater performance.

In comparison to other machine learning approaches for gene design, OrthologTransformer is unique. Most deep-learning codon optimizers – including ICOR^29^, CodonBERT^30^ and CodonTransformer^24^ – frame expression tuning as a synonym-only sequence-to-sequence task that leaves the amino-acid chain untouched. Those models can capture complex codon usage features and even context-dependent effects on gene expression, yet they still operate within the traditional “no amino-acid change” paradigm originally popularised by frequency-based codon-optimization tools such as JCat^31^. OrthologTransformer expands this paradigm and can be seen as bridging evolutionary bioinformatics and synthetic biology: it uses evolution’s imprint (the variations between orthologs^32^) to guide synthetic gene construction. In doing so, it creates a new category of tool that we might call an “ortholog designer.” By incorporating insertions and deletions, OrthologTransformer even allows changes in protein length if that has been characteristic for those genes between species.

Recent work shows that co-optimising multiple constraints can greatly enhance mRNA performance: one algorithm that jointly tunes folding energy and codon usage raised vaccine antibody titres 128-fold^33^, and another design strategy used measured half-life data to increase both expression and decay resistance^34^. Building on this insight, we feed OrthologTransformer’s generative sequences into a Monte-Carlo Tree Search that simultaneously balances GC content and folding energy, yielding designs that satisfy host adaptation, synthesis feasibility and translational robustness in a single step.

The superior performance of the AI9-designed PETase in *B. subtilis* also has environmental implications. PET plastic accumulation is a global concern, and biodegradation is one approach to address it^35,36^. While *I. sakaiensis* itself can degrade PET, it grows slowly and is not ideal for large-scale applications. *B. subtilis*, on the other hand, is a fast-growing, spore-forming bacterium amenable to fermentation^37^, and has been increasingly recognized as a host for heterologous PETase expression^38–40^. By empowering *B. subtilis* to produce active PETase, we move closer to a system that could be used to biodegrade or upcycle PET waste. Notably, *B. subtilis* secretes the enzyme, potentially allowing continuous enzyme action on plastic without cell lysis steps. Our SEM images showing *B. subtilis* physically boring into plastic are a vivid demonstration of how a harmless soil bacterium can be reprogrammed to attack a synthetic polymer. This showcases the promise of synthetic biology for environmental remediation – with AI helping to surmount the expression barriers that have historically limited such efforts. Although OrthologTransformer dramatically enlarges the accessible design space, three hurdles still limit its reliability. First, seven PETase variants (AI1–AI7) show mean pLDDT values below 70, a confidence band enriched in intrinsically disordered segments and typically > 3 Å RMSD from experimental structures^41,42^; adding an explicit 3-D objective during generation might therefore improve foldability. Second, the model does not actively preserve catalytic or binding motifs; both enzyme-design case studies and solubility-focused design efforts have documented apparently well-folded or stable sequences that nonetheless lose activity because of subtle active-site shifts^43,44^. Finally, OrthologTransformer is currently trained and tested exclusively on bacterial sequences. As such, it does not yet support design tasks involving eukaryotic genes, which often involve more complex regulation such as splicing, compartmental localization, or post-translational modifications. Extending its applicability to eukaryotic systems would represent a major step toward broader utility in synthetic biology. Integrating a rapid structure-confidence filter, constraining functional residues, and expanding cross-domain training data should together raise the in-cell success rate of future OrthologTransformer designs.

## Conclusion

We have demonstrated that OrthologTransformer enables a comprehensive approach to gene retargeting across species, encompassing synonymous codon optimization and intelligent non-synonymous modifications. By training on nature’s own solutions to cross-species adaptation (orthologs), the model learns to balance the dual demands of maintaining protein function and adapting to a new genomic context. The successful prediction and experimental validation of a functional PETase ortholog exemplifies the power of this approach. OrthologTransformer thus differentiates itself from existing methods as a tool that doesn’t just optimize codon usage – it effectively engineers orthologous genes, bringing us closer to the longstanding goal of seamless genetic feature transfer between organisms. We anticipate that this capability will find broad utility in the synthetic biology community and beyond, enabling innovations from improved bioindustrial enzyme production to novel therapeutics and environmental solutions.

## Material and Methods

### Datasets Construction

Development of the OrthologTransformer model required large-scale, high-quality ortholog datasets to accurately capture diverse genetic variations, including synonymous and non-synonymous substitutions, as well as insertions and deletions (indels). A strict separation between training and test datasets was also essential to rigorously evaluate model generalization. Initially, we processed sequence data from 2 to 132 bacterial species retrieved from RefSeq using OrthoFinder^45^. To ensure high data quality, sequences were restricted to lengths under 2,100 bp, and only ortholog pairs with length ratios between 0.97 and 1.03 were selected. For constructing a larger and more comprehensive dataset, we leveraged the OMA database, which provides highly reliable ortholog data identified through rigorous algorithms. Distinct orthologous groups were allocated separately to training and testing sets, enabling accurate evaluation of the model’s predictive versatility. The finalized training dataset comprised 1,071,976 orthologous groups and 4,971,060 ortholog pairs derived from 2,138 bacterial species. To further enhance dataset quality, we applied gap-processing by removing alignment gaps following amino acid alignments, thus preserving conservation in functionally important regions. Both gap-processed and unprocessed datasets were prepared to maintain a balance between evolutionary information retention and alignment quality. For the test set, we deliberately selected species pairs with diverse GC contents and optimal growth temperatures to examine how biological characteristics impact conversion accuracy. This multifaceted approach enables OrthologTransformer to achieve robust and highly accurate predictions even for genes it has not encountered previously.

### OrthologTransformer Architecture

We present OrthologTransformer, a Sequence-to-Sequence model specifically designed for cross-species DNA sequence conversion (Figure 1). The model employs a Transformer architecture with a critical innovation: species signal tokenization. Species tokens (*s*_*src*_for source and *s*_*tgt*_ for target) are prepended to encoder and decoder inputs respectively, enabling explicit indication of conversion direction. The architecture comprises 20-layer encoder and decoder stacks with 114,095,940 total parameters. DNA sequences are tokenized at codon resolution, with each codon and species signal mapped to 512-dimensional trainable embeddings. We employ 16-head multi-head attention mechanisms and position-wise feed-forward networks in each layer. The model utilizes cross-entropy loss with Adam optimizer, operating with the following hyperparameters: embedding dimension 512, feed-forward dimension 1024, dropout rate 0.0, batch size 64, and initial learning rate 0.0001. During inference, the decoder generates sequences autoregressively, selecting the most probable codon at each step from a 67-dimensional softmax output.

### Two-stage Learning Strategy

Our training approach combines pretraining on diverse species relationships with targeted fine-tuning (Figure 1b). During pretraining, the model learns general sequence conversion patterns through many-to-many converting tasks across all available species pairs. In the fine-tuning phase, we specialize the model for specific one-to-one species pair conversions, such as *I. sakaiensis* to *B. subtilis*, using only relevant ortholog pairs to capture species-specific sequence features. This two-stage strategy enables OrthologTransformer to achieve both general sequence conversion capabilities and species-specific optimization.

### PETase Sequence Design

To generate *B. subtilis*-compatible PETase sequences from *I. sakaiensis*, we strategically initialized the decoder with the target species token *s*_*tgt*_ concatenated with a codon-optimized signal sequence derived from the original PETase. This initialization guides the model toward generating sequences suitable for expression in the target organism. We implemented a comprehensive design strategy that explored multiple dimensions of sequence generation to create a diverse set of PETase variants (Table 3). Our approach systematically varied several key parameters: the choice of signal sequence (either codon-optimized or AI-designed), whether training data underwent alignment preprocessing, the source database (OrthoFinder or OMA) with corresponding variation in species diversity (ranging from 23 to 2,138 species), the application of MCTS for sequence optimization, and whether the model underwent fine-tuning for the specific species pair. This multifaceted approach allowed us to generate twelve distinct sequences that collectively represent different combinations of biological knowledge integration, computational optimization, and AI-driven design, enabling us to evaluate which factors most significantly influence successful heterologous expression.

### Monte Carlo Tree Search Optimization

We developed a MCTS algorithm to optimize generated sequences for both GC content and RNA secondary structure stability. This addresses the competing requirements of synthetic accessibility (favoring low GC content around 0.36) and functional mRNA stability (requiring stable secondary structures). Specifically, while lowering GC content facilitates DNA synthesis and cloning, maximizing translational output requires mRNA secondary structures stabilized to an optimal degree, extending transcript half-life without hindering ribosomal access^34,46^. Our MCTS implementation builds a search tree where each node represents a partial DNA sequence, with child nodes adding additional codons based on Transformer-generated probability distributions. We select the top three most probable tokens for expansion at each step. Node evaluation employs a reward function that balances GC content optimization with secondary structure stability:

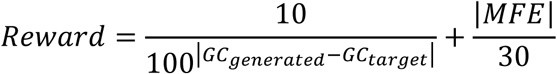

where *GC*_*generated*_ represents the sequence’s GC content, *GC*_*target*_ equals 0.36, and *MFE* denotes minimum free energy of the RNA structure. The algorithm iteratively performs selection using UCB1^47^, expansion based on model probabilities, simulation to terminal states, and backpropagation of rewards, enabling efficient exploration of the sequence space while maintaining biological constraints.

### Evaluation Metrics

To comprehensively evaluate the sequences generated by our approach, we employed multiple metrics spanning sequence identity, structural prediction, and biological compatibility. For sequence evaluation, we calculated codon sequence identity by performing pairwise alignment using the Needleman-Wunsch algorithm with match score 1 and gap penalty 0, followed by normalization against sequence length. Similarly, we computed amino acid sequence similarity by pairwise alignment based on the BLOSUM score, applying gap penalties of 10 for opening and 0.1 for extension, with scores normalized by sequence length. For structural assessment, we utilized AlphaFold3 to compute TM-scores comparing predicted structures of AI-designed PETase variants against the wild-type PETase structure, providing quantitative evaluation of structural conservation. We also analyzed GC content, a critical parameter affecting both synthetic accessibility and thermal stability, ensuring compatibility with the target organism’s genomic characteristics. Translation efficiency was evaluated using the Codon Adaptation Index (CAI), calculated as:

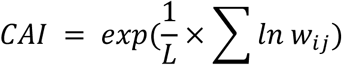

where *L* represents gene length in codons, and *w*_*ij*_ denotes the relative adaptiveness of each codon, defined as:

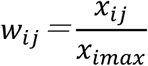

Here, *x*_*ij*_ represents the frequency of synonymous codon *j* for amino acid *i*, while *x*_*imax*_ indicates the maximum frequency among synonymous codons for the same amino acid. We computed CAI values using whole-genome codon usage frequencies as reference data for each species, with values ranging from 0 to 1, where values closer to 1 indicate optimal codon usage for the target organism.

### Experimental Procedures

#### Strain construction

Synthesized genes and the expression vector (*E. coli*/*B. subtilis* shuttle vector pUB980-2^48^) were amplified by PCR. The resulting DNA fragments were mixed at a 1:1 molar ratio (vector:gene) and assembled using NEBuilder HiFi DNA Assembly Master Mix (NEB, Japan). After incubation at 50°C for 30 minutes, the assembled DNA was transformed into *E. coli* HST08 (Takara, Japan), and plasmids were subsequently extracted. These plasmids were then transformed into *B. subtilis* RM125^49^ using the method described by Spizizen^50^. The strains and plasmids used or constructed in this study are listed in Supplementary Table 1 and 2 (Supplementary Materials and Methods).

#### RNA extraction and reverse transcription

Recombinant *Bacillus subtilis* cells were harvested during the logarithmic growth phase (Supplementary Figure MT_2) by centrifugation at 8,000 × g for 3 minutes (refer to the Supplementary Materials and Methods for details). Three biological replicates were prepared for each recombinant strain. The cells were resuspended in 100 μL of a 5 mg/mL lysozyme solution prepared in 1× TE buffer and incubated at 37°C for 10 minutes. RNA extraction was performed using the NucleoSpin RNA kit (Macherey-Nagel, Germany) according to the manufacturer’s instructions. Total RNA was quantified and assessed for quality using the RNA 6000 Nano Kit on an Agilent 2100 Bioanalyzer (Agilent Technologies, USA). All samples exhibited an RNA integrity number (RIN) greater than 9.8. Reverse transcription of total RNA was performed using the ReverTra Ace qPCR RT Master Mix with gDNA Remover (Toyobo Co., Ltd., Japan) according to the manufacturer’s guidelines.

#### PCR for full-length transcript

PCR was then performed using a forward primer binding near the 5’ end of the PETase coding sequence (within the signal peptide region) and a reverse primer at the 3’ end (within the His-tag region). Amplification was performed using PrimeSTAR Max DNA Polymerase Ver.2 (Takara, Japan), with the following thermal cycling conditions: 20 cycles of 98°C for 10 seconds, 55°C for 10 seconds, and 68°C for 5 seconds. The expected amplicon was visualized on agarose gel. All PETase constructs yielded the correct band from cDNA, confirming presence of full-length mRNA.

#### Quantitative real-time PCR (qPCR)

For quantification of PETase transcripts, we performed qPCR on a StepOne Plus Real-Time PCR System (Thermo Fisher Scientific, USA) with Power SYBR Green PCR Master Mix (Applied Biosystems, USA). Gene-specific primers were designed to amplify an internal region within the signal sequence of PETase gene, which is common to all recombinant strains. As an internal control, primers for the single-copy *gyrA* gene of *B. subtilis* (DNA gyrase subunit A) were used. The ΔCt for PETase was calculated relative to gyrA, and then expression levels were normalized to the WT (wild-type) PETase sample (set as 1.0) using the ΔΔCt method.

#### Western blotting (immunoblot)

To detect PETase protein, cultures (3 mL in LB + kanamycin + xylose) were grown for 12 h, and samples of both the culture supernatant and cell pellet were collected. For the supernatant, 1 mL of culture was centrifuged (8,000 rpm, 5 min, 4℃) to remove cells, and the supernatant was mixed with 6x sample loading buffer. All samples were normalized by culture volume (since cell density was similar across strains at 24 h), and boiled (5 min, 95℃) before SDS-PAGE. Proteins were separated on 12.5% SDS-PAGE gels. For secreted proteins, we loaded 9.0 µL of culture supernatant samples directly. After electrophoresis, proteins were transferred to a PVDF membrane. The membrane was probed with an anti-polyhistidine monoclonal antibody (against the 6×His tag at PETase C-terminus) at 1:5000 dilution, followed by an Alkaline Phosphate-conjugated secondary antibody. Detection was performed with BCIP/NBT and recorded on a imager. A His-tagged protein ladder was used to verify molecular weight.

### PETase Activity Assessment

#### PET film degradation experiment

Additive-free PET film (thickness ∼0.25 mm) was cut into ∼2 cm² pieces (4 cm × 5 cm rectangles). Before use, the films were thoroughly washed with 1% (w/v) sodium dodecyl sulfate (SDS) for 30 min and rinsed with Milli-Q water, to remove any contaminants and surface treatments. Films were then soaked in 70% ethanol for 1 h for sterilization and air-dried in a laminar flow hood. Each PET film piece was placed in a 100 mL Erlenmeyer flask containing 20 mL of LB medium with kanamycin (25 µg/mL). *B. subtilis* transformant pre-cultures were grown as described above, and 200 µL of the overnight culture was used to inoculate the flask (initial OD ∼0.1). 1% xylose was added at inoculation to induce PETase expression from the start. Cultures with PET film were incubated at 30 °C, 180 rpm for up to 7 days. Flasks were covered but not sealed (to allow aeration). At predetermined time points (24 h, 48 h, 72 h, and 7 days), 0.5 mL of culture medium was sampled for HPLC analysis of PET degradation products, using the same protocol as for BHET samples. Because PET is solid and only its soluble hydrolysis products are in the medium, we focused on MHET and TPA detection. BHET was not expected unless PET hydrolysis stops at dimer stage (which PETase typically does not). Each assay was done in triplicate flasks for each strain. After 7 days, the PET film was recovered from the culture for imaging.

#### HPLC quantification of PET hydrolysis products

Culture supernatants from PET film flasks were processed as for BHET samples (acetonitrile/formic acid quench and centrifugation). The HPLC analysis was identical. We confirmed linearity of MHET detection in the relevant range (0–50 µM) by running standard solutions. In control flasks (no PETase), no MHET or TPA peaks were observed at any time point. In PETase-expressing samples, a clear MHET peak emerged by day 2–3. MHET concentration was quantified for each sample. TPA was not observed (likely remaining bound in undetectable form or not produced due to lack of MHETase). The results were plotted as MHET (µmol) released per flask over time. Statistical comparison of MHET at day 3 among strains was performed by one-way ANOVA with Tukey’s post-hoc test. AI-12 produced significantly more MHET than all other strains (***p<0.001 for AI-12 vs each other PETase strain). These quantitative data corroborate the qualitative ranking of enzyme performance.

#### Scanning Electron Microscopy (SEM)

After 7 days of incubation, PET film pieces were retrieved from the cultures with sterile tweezers. To remove attached cells and biofilm, each film was gently sonicated in 2% SDS solution for 5 min, then rinsed thoroughly with Milli-Q water (several dips). Films were then air-dried completely. The dried films were mounted on SEM stubs and sputter-coated with a thin layer of osmium (5 nm thickness) using an osmium plasma coater (HPC-20, Vacuum Device). Coating with osmium avoids charging and gives high-resolution imaging of polymer surfaces. The samples were examined under a field-emission scanning electron microscope (FEI Inspect S50) at an accelerating voltage of 5 kV. Images were taken at 1000× to 10,000× magnifications. We focused on 10,000× for detailed surface features. Multiple fields were imaged for each film. The control film showed a smooth surface with machining striations but no biological etching. Treated films showed various degrees of erosion: we observed pits, grooves, and in some cases, large holes that appear to correspond to where a bacterial cell was attached (as inferred by size and shape). Representative images for empty vector (no PETase) vs. AI-12 vs. AI-9 were presented in Fig. 8C–E. As reported in Results, AI-12’s film had ∼1 µm holes and extensive surface roughening, whereas AI-9 had fewer and smaller pits, and the control had none. These SEM results visually confirm PET hydrolysis and were consistent across triplicate samples.

## Code Availability

The source code will be released upon the official publication of the paper in a peer-reviewed journal.

## Declarations

### Authors’ Contributions

MA; implemented the software, analysed data, and compared with the existing methods. MT; performed strain constructions and western blotting analysis. YH, KM; performed PET degradation activity assay. MU; performed mRNA full-length transcription and qPCR analysis. MI; designed strain constructions. TK, MK; performed western blotting analysis. YS; designed and supervised the research, analysed data. MA, MT, YH, MU, and YS wrote and edited the paper. All authors read and approved the final manuscript.

## Acknowledgements

This work was supported by JST, CREST Grant Number JPMJCR20S3, Japan.

## References

1. Gustafsson, C., Govindarajan, S. & Minshull, J. Codon bias and heterologous protein expression. Trends Biotechnol. 22, 346–353 (2004).

2. Plotkin, J. B. & Kudla, G. Synonymous but not the same: the causes and consequences of codon bias. Nat. Rev. Genet. 12, 32–42 (2011).

3. Sharp, P. M. & Li, W. H. The codon Adaptation Index--a measure of directional synonymous codon usage bias, and its potential applications. Nucleic Acids Res. 15, 1281– 1295 (1987).

4. Ranaghan, M. J., Li, J. J., Laprise, D. M. & Garvie, C. W. Assessing optimal: inequalities in codon optimization algorithms. BMC Biol. 19, 36 (2021).

5. Paremskaia, A. I. et al. Codon-optimization in gene therapy: promises, prospects and challenges. Front. Bioeng. Biotechnol. 12, 1371596 (2024).

6. Qian, W., Yang, J.-R., Pearson, N. M., Maclean, C. & Zhang, J. Balanced codon usage optimizes eukaryotic translational efficiency. PLoS Genet. 8, e1002603 (2012).

7. Mauro, V. P. Codon optimization in the production of recombinant biotherapeutics: Potential risks and considerations. BioDrugs 32, 69–81 (2018).

8. Boël, G. et al. Codon influence on protein expression in E. coli correlates with mRNA levels. Nature 529, 358–363 (2016).

9. Tuller, T., Waldman, Y. Y., Kupiec, M. & Ruppin, E. Translation efficiency is determined by both codon bias and folding energy. Proc. Natl. Acad. Sci. U. S. A. 107, 3645–3650 (2010).

10. Presnyak, V. et al. Codon optimality is a major determinant of mRNA stability. Cell 160, 1111–1124 (2015).

11. Fitch, W. M. Distinguishing homologous from analogous proteins. Syst. Zool. 19, 99–113 (1970).

12. Escorcia-Rodríguez, J. M., Esposito, M., Freyre-González, J. A. & Moreno-Hagelsieb, G. Non-synonymous to synonymous substitutions suggest that orthologs tend to keep their functions, while paralogs are a source of functional novelty. PeerJ 10, e13843 (2022).

13. Goodman, D. B., Church, G. M. & Kosuri, S. Causes and effects of N-terminal codon bias in bacterial genes. Science 342, 475–479 (2013).

14. Angov, E., Hillier, C. J., Kincaid, R. L. & Lyon, J. A. Heterologous protein expression is enhanced by harmonizing the codon usage frequencies of the target gene with those of the expression host. PLoS One 3, e2189 (2008).

15. Kudla, G., Murray, A. W., Tollervey, D. & Plotkin, J. B. Coding-sequence determinants of gene expression in Escherichia coli. Science 324, 255–258 (2009).

16. Buhr, F. et al. Synonymous codons direct cotranslational folding toward different protein conformations. Mol. Cell 61, 341–351 (2016).

17. Fu, H. et al. Codon optimization with deep learning to enhance protein expression. Sci. Rep. 10, 17617 (2020).

18. Outeiral, C. & Deane, C. M. Codon language embeddings provide strong signals for use in protein engineering. Nature Machine Intelligence 6, 170–179 (2024).

19. Tunney, R. et al. Accurate design of translational output by a neural network model of ribosome distribution. Nat. Struct. Mol. Biol. 25, 577–582 (2018).

20. Yoshida, S. et al. A bacterium that degrades and assimilates poly(ethylene terephthalate). Science 351, 1196–1199 (2016).

21. Vaswani, A. et al. Attention is all you need. in Advances in neural information processing systems 5998–6008 (papers.nips.cc, 2017).

22. Sutskever, I., Vinyals, O. & Le, Q. V. Sequence to sequence learning with Neural Networks. Neural Inf Process Syst abs/1409.3215, 3104–3112 (2014).

23. Altenhoff, A. M. et al. OMA orthology in 2024: improved prokaryote coverage, ancestral and extant GO enrichment, a revamped synteny viewer and more in the OMA Ecosystem. Nucleic Acids Res. 52, D513–D521 (2024).

24. Fallahpour, A., Gureghian, V., Filion, G. J., Lindner, A. B. & Pandi, A. CodonTransformer: a multispecies codon optimizer using context-aware neural networks. Nat. Commun. 16, 3205 (2025).

25. Joo, S. et al. Structural insight into molecular mechanism of poly(ethylene terephthalate) degradation. Nat. Commun. 9, 382 (2018).

26. Tenthorey, J. L. et al. Indels allow antiviral proteins to evolve functional novelty inaccessible by missense mutations. Cell Genom. 100818 (2025).

27. Tournier, V. et al. An engineered PET depolymerase to break down and recycle plastic bottles. Nature 580, 216–219 (2020).

28. Austin, H. P. et al. Characterization and engineering of a plastic-degrading aromatic polyesterase. Proc. Natl. Acad. Sci. U. S. A. 115, E4350–E4357 (2018).

29. Jain, R., Jain, A., Mauro, E., LeShane, K. & Densmore, D. ICOR: improving codon optimization with recurrent neural networks. BMC Bioinformatics 24, 132 (2023).

30. Ren, Z. et al. CodonBERT: a BERT-based architecture tailored for codon optimization using the cross-attention mechanism. Bioinformatics 40, btae330 (2024).

31. Grote, A. et al. JCat: a novel tool to adapt codon usage of a target gene to its potential expression host. Nucleic Acids Res. 33, W526–31 (2005).

32. Koonin, E. V. Orthologs, paralogs, and evolutionary genomics. Annu. Rev. Genet. 39, 309–338 (2005).

33. Zhang, H. et al. Algorithm for optimized mRNA design improves stability and immunogenicity. Nature 621, 396–403 (2023).

34. Leppek, K. et al. Combinatorial optimization of mRNA structure, stability, and translation for RNA-based therapeutics. Nat. Commun. 13, 1536 (2022).

35. Wei, R. & Zimmermann, W. Microbial enzymes for the recycling of recalcitrant petroleum-based plastics: how far are we? Microb. Biotechnol. 10, 1308–1322 (2017).

36. Geyer, R., Jambeck, J. R. & Law, K. L. Production, use, and fate of all plastics ever made. Sci. Adv. 3, e1700782 (2017).

37. Wang, Z. et al. Bacillus subtilis as an excellent microbial treatment agent for environmental pollution: A review. Biotechnol. J. 20, e70026 (2025).

38. Wang, N. et al. Enhancing secretion of polyethylene terephthalate hydrolase PETase in Bacillus subtilis WB600 mediated by the SPamy signal peptide. Lett. Appl. Microbiol. 71, 235–241 (2020).

39. Qi, X., Yan, W., Cao, Z., Ding, M. & Yuan, Y. Current advances in the biodegradation and bioconversion of polyethylene terephthalate. Microorganisms 10, 39 (2021).

40. Huang, X. et al. Tat-independent secretion of polyethylene terephthalate hydrolase PETase in Bacillus subtilis 168 mediated by its native signal peptide. J. Agric. Food Chem. 66, 13217–13227 (2018).

41. Tunyasuvunakool, K. et al. Highly accurate protein structure prediction for the human proteome. Nature 596, 590–596 (2021).

42. Alderson, T. R., Pritišanac, I., Kolarić, Đ., Moses, A. M. & Forman-Kay, J. D. Systematic identification of conditionally folded intrinsically disordered regions by AlphaFold2. Proc. Natl. Acad. Sci. U. S. A. 120, e2304302120 (2023).

43. Khersonsky, O. & Fleishman, S. J. What have we learned from design of function in large proteins? Biodes. Res. 2022, 9787581 (2022).

44. Qing, R. et al. Protein design: From the aspect of water solubility and stability. Chem. Rev. 122, 14085–14179 (2022).

45. Emms, D. M. & Kelly, S. OrthoFinder: phylogenetic orthology inference for comparative genomics. Genome Biol. 20, 238 (2019).

46. Noskov, V. N. et al. Assembly of large, high G+C bacterial DNA fragments in yeast. ACS Synth. Biol. 1, 267–273 (2012).

47. Coulom, R. Efficient selectivity and backup operators in Monte-Carlo tree search. In Computers and Games 72–83 (2007).

48. Zhao, X. et al. High copy number and highly stable Escherichia coli-Bacillus subtilis shuttle plasmids based on pWB980. Microb. Cell Fact. 19, 25 (2020).

49. Uozumi, T. et al. Restriction and modification in Bacillus species: genetic transformation of bacteria with DNA from different species, part I. Mol. Gen. Genet. 152, 65–69 (1977).

50. Spizizen, J. Transformation of biochemically deficient strains of Bacillus subtilis by deoxyribonucleate. Proc. Natl. Acad. Sci. U. S. A. 44, 1072–1078 (1958).

