## Supplementary Materials for "Generative AI-driven artificial DNA design for enhancing inter-species gene activation and enzymatic degradation of PET"

Supplemental Methods

### Monte Carlo Tree Search for Multi-objective Sequence Optimization

Monte Carlo Tree Search (MCTS) was employed for sequence optimization in this study, particularly to adapt the PETase enzyme from Ideonella sakaiensis for optimal expression in Bacillus subtilis. DNA sequence design requires balancing multiple objectives, with GC content and RNA secondary structure stability playing critical roles.

GC content represents the proportion of guanine (G) and cytosine (C) bases in DNA. Lower GC content can facilitate chemical synthesis, enabling production of longer sequences [S1]. However, this creates a trade-off, as reduced GC content tends to decrease RNA secondary structure stability, which is essential for RNA functionality and expression efficiency.

Our approach aimed to achieve a GC content close to 0.36 while maximizing RNA secondary structure stability. To navigate this multi-objective optimization problem, we implemented MCTS with a reward function that balanced both objectives:

$$Reward = \frac{10}{100^{|GC_{generated} - GC_{target}|}} + \frac{|MFE|}{30}$$

Where $GC_{generated}$ represents the GC content of the designed sequence, $GC_{target}$ equals 0.36, and MFE denotes the minimum free energy of the RNA structure. The algorithm proceeded through four key steps:

Selection: Choosing optimal nodes using the UCB1 algorithm, calculated as:

$$UCB1 = \frac{w_i}{n_i} + 2\sqrt{\frac{ln\ N}{n_i}}$$

where $w_i$ is the cumulative reward of node $i$, $n_i$ is the visit count of node $i$, and $N$ is the visit count of the parent node.

Expansion: Generating child nodes based on the top three probabilities from the Transformer model

Simulation: Calculating rewards by evaluating GC content and structure stability

Backpropagation: Updating cumulative rewards and visit counts from selected nodes to the root

Figure S.3. tracks the optimization process for the AI9 PETase sequence across four key metrics: reward value, GC content, MFE, and tree depth. Our analysis revealed a two-phase optimization pattern where MCTS rapidly improved rewards from 15 to 18 during the first 200,000 iterations, followed by a gradual increase to 22 by iteration 300,000 (Figure S.3.a). This pattern reflects the algorithm's initial focus on GC content optimization before transitioning to RNA structure refinement.

The algorithm effectively converged toward the target GC content of 0.36 (Figure S.3.b), while simultaneously optimizing RNA secondary structure. The minimum free energy (MFE) showed a characteristic progression from an initial -220 kcal/mol to approximately -245 kcal/mol during the first 200,000 iterations, eventually stabilizing around -280 kcal/mol (Figure S.3.c).

Notably, our MCTS implementation successfully navigated the inherent trade-off between GC content and RNA stability. As shown in Figure S.3.b and c, AI9 achieved both a low GC content of 0.37 and a highly stable RNA structure with an MFE of -281 kcal/mol. This resolution of competing objectives highlights the efficacy of MCTS for complex biological sequence design problems, aligning with recent research demonstrating MCTS effectiveness for multi-objective optimization in biological contexts [S2]. Furthermore, Kathrin et al. [S3] have shown that optimizing solely for CAI does not necessarily improve translation efficiency or stability; rather, balancing structure stability and translation efficiency is critical.

**Plasmids information**

The *E. coli*/*B. subtilis* shuttle vector pUB980-2-$P_{xyl}$, a derivative of pUB980-2 [S4], was used as the expression vector in this study. This plasmid carries a kanamycin resistance gene functional in both *E. coli* and *B. subtilis*, with working concentrations of 50 µg/mL and 25 μg/mL, respectively. Gene expression is driven by the $P_{xyl}$ promoter (5'-CTAAAAAAAATATTGAAAATACTGACGAGGTTATATAAGATGAAAATAAGTT AGTTTGTTTAAACAACAAACTAATAGGTGA-3') in the presence of xylose, and a 6xHis-tag allows for detection of the expressed protein. For extracellular secretion of PETase, the signal peptide $SP_{sacC}$ (sequence: MKKRLIQVMIMFTLLLTMAFSADA), derived from *B. subtilis*, was employed. All plasmids constructed and used in this study are listed in Table S.1 and S.2.

**RNA extraction and reverse transcription**

RNA extraction and reverse transcription: To assess the transcription of the PETase gene, recombinant B. subtilis strains were cultured overnight in LB medium supplemented with kanamycin (25 μg/mL). A portion of the overnight culture was diluted into fresh LB medium to an initial optical density at 600 nm ($OD_{600}$) of 0.1. Cultures were incubated at 37°C with shaking, and PETase expression was induced by adding 1% xylose at the time of inoculation. Recombinant B. subtilis cells were harvested during the logarithmic growth phase by centrifugation at 8,000 × g for 3 minutes. For each recombinant strain, three biological replicates were prepared. Genetically engineered B. subtilis strains were centrifuged at 8,000 g for 3 minutes during the logarithmic growth phase, and samples were collected for three replicates per recombinant strain. The cells were suspended in 100 μL of a 5 mg/mL lysozyme solution prepared in 1× TE buffer and incubated at 37°C for 10 minutes. RNA extraction was carried out using the Nucleo Spin RNA kit (Macherey-Nagel, Germany), following the manufacturer's instructions. Total RNA was quantified and assessed for quality using the RNA 6000 Nano Kit on an Agilent 2100 Bioanalyzer (Agilent Technologies, USA). All samples exhibited an RNA integrity number (RIN) greater than 9.8. Total RNA was reverse transcribed using the ReverTra Ace qPCR RT Master Mix with gDNA Remover (Toyobo Co., Ltd., Japan), according to the manufacturer’s guidelines. The reverse transcription reaction included RNA denaturation, gDNA removal, and synthesis of cDNA, with an initial total RNA input of 500 ng for each 10 μL of cDNA solution.

Supplemental Tables

**Supplementary Table 1.** Plasmids used in this study.

| **plasmids** | **relevant properties** |
|---|---|
| pUB980-2 | [S4] |
| pUB980-2-$P_{xyl}$ (EV) | Derivative of pUB980-2.<br>$Km^r$, $Km^r$, $P_{xyl}$ promoter,<br>*E. coli/B. subtilis* shuttle vector |
| pUB980-2- $P_{xyl}$ -$SP_{sacC}$-WT | pUB980-2-$P_{xyl}$ derivative,<br>$SP_{sacC}$-WT |
| pUB980-2- $P_{xyl}$ -$SP_{sacC}$-CO | pUB980-2-$P_{xyl}$ derivative,<br>$SP_{sacC}$-CO |
| pUB980-2- $P_{xyl}$ -$SP_{sacC}$-AI1 | pUB980-2-$P_{xyl}$ derivative,<br>$SP_{sacC}$-AI1 |
| pUB980-2- $P_{xyl}$ -$SP_{sacC}$-AI2 | pUB980-2-$P_{xyl}$ derivative,<br>$SP_{sacC}$-AI2 |
| pUB980-2- $P_{xyl}$ -$SP_{sacC}$-AI3 | pUB980-2-$P_{xyl}$ derivative,<br>$SP_{sacC}$-AI3 |
| pUB980-2- $P_{xyl}$ -$SP_{sacC}$-AI4 | pUB980-2-$P_{xyl}$ derivative,<br>$SP_{sacC}$-AI4 |
| pUB980-2- $P_{xyl}$ -$SP_{sacC}$-AI5 | pUB980-2-$P_{xyl}$ derivative,<br>$SP_{sacC}$-AI5 |
| pUB980-2- $P_{xyl}$ -$SP_{sacC}$-AI6 | pUB980-2-$P_{xyl}$ derivative,<br>$SP_{sacC}$-AI6 |
| pUB980-2- $P_{xyl}$ -$SP_{sacC}$-AI7 | pUB980-2-$P_{xyl}$ derivative,<br>$SP_{sacC}$-AI7 |
| pUB980-2- $P_{xyl}$ -$SP_{sacC}$-AI8 | pUB980-2-$P_{xyl}$ derivative,<br>$SP_{sacC}$-AI8 |
| pUB980-2- $P_{xyl}$ -$SP_{sacC}$-AI9 | pUB980-2-$P_{xyl}$ derivative,<br>$SP_{sacC}$-AI9 |

| | |
|---|---|
| pUB980-2- $P_{xyl}$ -$SP_{sacC}$-AI10 | pUB980-2-$P_{xyl}$ derivative,<br>$SP_{sacC}$-AI10 |
| pUB980-2- $P_{xyl}$ -$SP_{sacC}$-AI11 | pUB980-2-$P_{xyl}$ derivative,<br>$SP_{sacC}$-AI11 |
| pUB980-2- $P_{xyl}$ -$SP_{sacC}$-AI12 | pUB980-2-$P_{xyl}$ derivative,<br>$SP_{sacC}$-AI12 |

**Supplementary Table 2.** Strains used in this study.

| strains | relevant properties |
|---|---|
| *Escherichia coli* HST08 | Purchased from TaKaRa (Japan), *F－, endA1, supE44, thi-1, recA1, relA1, gyrA96, phoA, Φ80dlacZΔM15, Δ(lacZYA-argF) U169, Δ(mrr-hsdRMS-mcrBC), ΔmcrA, λ－* |
| *Bacillus subtilis* RM125 | [S5] *argA15, leuA8, SP90(s), hsdR168, hsdM168* |
| RM125-WT | *B. subtilis* RM125 containing pUB980-2-$P_{xyl}$-$SP_{sacC}$-WT |
| RM125-CO | *B. subtilis* RM125 containing pUB980-2-$P_{xyl}$-$SP_{sacC}$-CO |
| RM125-AI1 | *B. subtilis* RM125 containing pUB980-2-$P_{xyl}$-$SP_{sacC}$-AI1 |
| RM125-AI2 | *B. subtilis* RM125 containing pUB980-2-$P_{xyl}$-$SP_{sacC}$-AI2 |
| RM125-AI3 | *B. subtilis* RM125 containing pUB980-2-$P_{xyl}$-$SP_{sacC}$-AI3 |
| RM125-AI4 | *B. subtilis* RM125 containing pUB980-2-$P_{xyl}$-$SP_{sacC}$-AI4 |
| RM125-AI5 | *B. subtilis* RM125 containing pUB980-2-$P_{xyl}$-$SP_{sacC}$-AI5 |
| RM125-AI6 | *B. subtilis* RM125 containing pUB980-2-$P_{xyl}$-$SP_{sacC}$-AI6 |
| RM125-AI7 | *B. subtilis* RM125 containing pUB980-2-$P_{xyl}$-$SP_{sacC}$-AI7 |
| RM125-AI8 | *B. subtilis* RM125 containing pUB980-2-$P_{xyl}$-$SP_{sacC}$-AI8 |

| | |
|---|---|
| RM125-AI9 | *B. subtilis* RM125 containing pUB980-2-$P_{xyl}$-$SP_{sacC}$-AI9 |
| RM125-AI10 | *B. subtilis* RM125 containing pUB980-2-$P_{xyl}$-$SP_{sacC}$-AI10 |
| RM125-AI11 | *B. subtilis* RM125 containing pUB980-2-$P_{xyl}$-$SP_{sacC}$-AI11 |
| RM125-AI12 | *B. subtilis* RM125 containing pUB980-2-$P_{xyl}$-$SP_{sacC}$-AI12 |

**Supplementary Table 3.** AI-designed PETase DNA Sequences (AI1-AI12)

| Variant ID | DNA Sequence (5' → 3') |
|---|---|
| AI1 | 5'-ATGAATTTTCCGAGAGCATCAAGACTGATGCAAGCAGCAGTTCTGGGCGGCCTGATGGCAGTTTCA<br>GCAGCAGCAACAGCAAATAAAAACAATGACAAATGGACGATTCAGCAGGATGACACAGGATACACGATT<br>GAATTTCATGACAGATCAGGCAACACAAACACAAATCCGAATTTTCATACAGACAATAAAAAGCAATC<br>AAACAATTAGGCTGGAACTATAAATATTTAGGATTGAACAATCATCAAACAATTACAATTGATACAAAC<br>TCAAAATTAAACAAACCGACATCAAAATCATTGAAAAAACAATTGAAAGAAATTAACGGCATTAACCAT<br>CAAAACAACAGCAATCCAATCTATGGAAAAGTAGATACAAACCAATTAAAAGTGCCTACATTCGTATTT<br>GCAGGAGAAAACGACAGCATTGCCGGAACAAACTACAGCAAACTGACAATTTATAAAGAATTAAAAGGA<br>AATAAAGTTGTCTATGTGGGCACAAATGCTGGAAGAACCACAAACAGCTTTGCCAATAAAGTCAAATCA<br>TGGATGAAACAGTTCATGGATGAAGATGCAAACTACAGCACATTTGCATTTGAAAATCCAAACAGCACG<br>AAAGTCAACAGCTTTAAAAAAGCGGGCTTGAAATAA-3’ |
| AI2 | 5'-ATGAATTTTCCGAGAGCATCAAGACTGATGCAAGCAGCAGTTCTGGGCGGCCTGATGGCAGTTTCA<br>GCAGCAGCAACAGCAAAAACAGATCCAAACAGCCGAGGAATTGACCCGAACGCCGCTGCCATTCAATCT<br>GAAGCCAGTGCATTTATCTGGAAAGCGTTTACCGTTTCAAAACCGAGCAGCGGTGGAGCTGCTTCTGCG<br>TATTATACCTCACTTGCCAGCAGCACTGTGGGAGGAATTTCAGTACTACCGGGCTATACAGGGCGGGAT<br>GGCAGCATTTCCTGGTGGGGAGATGATATTGTCTCACACCAATGGTTGCTGGCGCTAGCAAGCACCAAT<br>TCTACATTGGATAATCCGGGAGGGAGCTCTCAGCGTCATTTGGACGCTTTCAAACGTGTGCAGACATTG<br>AAAAATGTGAAGGGAAATCCGATCTTTGGTACCATTGATCCCTTTAAAGTGGGCATTGCAAACTGGCAG<br>ATGGGAAGCACTGGCACGATATTAAGCTCGGTGAATGATCCGGCCTTTAAAACACTGACTTCCATAGAC<br>ATGCAAGATGAACAGTCTGAACTGCGCCAACTTACCGGTGTTACGAGCTTGTATCAATTAGCGCTTGAT<br>GCCATCAGCACCTATGAGGGCAAATCAATCCCCTTTGTCGATGATAACTCAACGGGAGCGAAAGACTTT<br>CTCACAGCGCTGGGAAGCTCCTTTGCTTGCTATTCCTCAGCCAATGCAAAAGCCAGCCTGTTGGGCATG<br>AAAAACGTTTTTATTTCGAAACGCCTCTCCGATGGAAAGCTGGCCTATCGGGACATTACGACGGTTTCC<br>GACGGTGATACGAGAGCGTCTGATTTCAAGCAAGTGCAGTGCCGTTAA-3’ |
| AI3 | 5'-ATGAAAATCAGACGAGCAAGCCGACTTTTTCAACTACTCGTGTTTACAACACTATTGGCAATAGCC<br>GGGGGTTTCACAAAAAGCTCCGATCCGTTGGCGATTGACACCAATCCATACCCTGCTGCCATACAGGCA<br>TCAGCTGCGGCATTTGTGGCGGGCAGTTTTATCACTGCAAAGCCGAGCGGCATCGGAGCTGATACAATG<br>TATTCGCAAAAAGATCTCGGAGGAACCGTTCTCGGGCTTTCAGTTGTGCCGGGTTTTACAGCAAAGGTG<br>AATTCTATGAAGTATTGGGGCAAAAGCCTCGTAGCATATCAATGGGCGGTTGCCAGCCAAGCAGTAAGC<br>AGCACATTCGATAATCCGTCAGGCAGAGCATCACAATATACGAAGGCGCTTCCAGAAACATTAAGCAGC<br>GGAAATGTCGGCAGCAGCAGCAGCGCCGGCGGCATCGATACGATGAAGGTAGGCATTCAAGGATGGCAG<br>CTGACAGATCAAGCCTTTATTTTAACGGAGGTAAACGGAGTGTCGTTTAAAGATGCCGGGAATACGAGC<br>GCCTGGAAGCTTTCCTCTGCCTTTCAACAAGTTACGGTGCCGCTGACAATTGTCGTCACCGATAACAAT<br>TCAGCTTACAATACAAATGCAAAATCATTGGCCATTTACGATGATACGTCCAGGGCAGCGAAACGCTTT<br>CTCACATTACGCGGCGGAAGTTCCCACGCCTATTCCTCTGCGAACATGGAGGTCGGATGGATGGGATGG<br>GAGGATGTTGCCTTTACTGAACAATTCTTTGACTCTGACTATACCTATCTGATCATCAAAACATCTTCA<br>GATGACGCTACCCGTTTAACTGAATTCAAGCACAGCACCTGCCAGTAA-3’ |
| AI4 | 5'-ATGAATTTTCCGAGAGCATCAAGACTGATGCAAGCAGCAGTTCTGGGCGGCCTGATGGCAGTTTCA<br>GCAGCAGCAACAGCATCTCCGGAAAAACAAAAAGAGAAATCGCTAAAGGTTGTTGCTTCTGATGGAACA<br>ACAGTATACTTCGGAAGAACAGAGCGTTCTTTTACAACGCTGAGTAAAGATGTAAGCGCCAATCGTGAT<br>ATCAAAAAAAAAGAGCCAAGCATCAATGATGAAGAAGCGAAGCTTCCGGTGATTCTTGGCACCTTTATG<br>ATACCACAAAGTTCTATGGATGTGAACATTACCATTCCGGTTTGGTTTCCTCATGATCAAAAAGGTGTT<br>TTAAACACAGGCTCATTCCCTCGTTCTACAACGTTTTATTCTTGTAAGGTTGGTGGAGTTGCCTCCGTT<br>GCATTATTAAAAGGTACTGTACTTAACAACTGCAATGAAAGAGGCTGGGCTGCAGGAGTATCAATTGCG<br>TCTGGACATGATGATGAACTTAACGGTGAAGCGAAAAAATGCAATTTGGGAGCGATGGCTGTTCCGGTT<br>GCTACTACAGCTGTCCGTTATCAGGAAAAAGGAATTTCTGCAGCTAATAAAACTGCGGTTGTTGATGGT<br>GGAGCACATGACGTTAACTGGGATAAAGAAGTTGTTCCCGCAATTTCCAAAGCTGGTGAAGAAAAAGGA<br>TCTTCTGGTTGGCTCTTAGAAGCAGGCGGAATTCTCAAACCTGTAGCTGGTGAAAAAGGATATCAACAT<br>ACACGCAACAATATGAGAGCTGGCTGGGCTATTCCAACATCTAATAAAGGAATTTACCACAACTGGGGA<br>GCTAAAGTATGCGGCTTGGTTCCTACTAACTTCCTTGATGTGTGGCAATGGACAGCTGGCTCACCAAAC<br>ACAGGTCATCCAGGCATGGAAGAAGAAGGTACAAGCAAATAA-3’ |

| | |
|---|---|
| AI5 | 5'-ATGAATTTTCCGAGAGCATCAAGACTGATGCAAGCAGCAGTTCTGGGCGGCCTGATGGCAGTTTCA<br>GCAGCAGCAACAGCAGCTGCTACCAAGCTGGCTGCCGGGGATATCCCGGGGGCCATGAAGCTGGAGGTC<br>AGCGCAAACAATGTCAGCCTCTGGCTGTTCCGCCAAAATGATCCTAGCAGCTTCGGAGCGGATGCCATG<br>TACTATCCGACAAACAATGGAGGCACCTATGGCGGGATTGCCATTATTCCCGGCTATACAGCTAAGCTG<br>GCTTCTTTGAAATGGTGGAAAGATTATGAAGTTGCGCATCAATATGTCTGCTTAACGATTGACTTAAAC<br>AGCCTGATGGATGAAACGAGCCAGCTCGGCACACAAAAGTGTGCGGTGCTTCGGGAGGTTGCCTCCATG<br>AAGGGAAAAGTATCTACCCCACTATTCGGCACCATCAACAAAAATAAGTGGGGAATTGCAAGCTGGAAA<br>GTGTCTAACAAAGGTGACTTCTTAACGGCTGTCCATAAAAACAGCTTTAAAACAGGCTGGAATCCGGCT<br>TTCTGGGATCAAAACTCTATCTATGACGGAAAAACATTCCCTTGGTTCCTGACATCACTGGAGCACAGC<br>AAAGCGAAGCCGTCCTCAAATCTATTTAATACGGTTTATGACAGCATGAGCAATTCAAGCAAGAAGTTT<br>ATTGAAATCTGGGGAGCGTCGCACAACTGCGCCGAGGTTGGCACTGCTGATGAAGGCTTTATCAACGGG<br>AAATCCGTTTCAGTCTTTAATCGATTCTTCGAACTCGATAAAAAATACCATACATTCCCGTGCAGCACG<br>CCGGTCGATAAACGTGTTGCCAATTATTCTACAAAATACTGCCAGTAA-3’ |
| AI6 | 5'-ATGAATTTTCCGAGAGCATCAAGACTGATGCAAGCAGCAGTTCTGGGCGGCCTGATGGCAGTTTCA<br>GCAGCAGCAACAGCAGCTGCTGCCGAGTTCTTAAAGGGTGATAACAATGTGTCTGTCAAGATAGATGAA<br>ACGGCTGGTTCGTTTAAGCTTTTTATTTTCACAGTTTCAAATCCGAGTTATTTCAGTGTCGGACAATTG<br>TATTATAAAGATAAAGCAATCAGCAGCTTGGGTGTCATTGCCATTATTCCTGGTTATACAGCTAAGTAT<br>GCTGATTTAAAATACTGGAAAAAAAAAATTAAAGCATATCAATTTAACTATTTGGCAGTTGATACAAAC<br>AATTTAAATGACATTGCATATGAAAATAATCTAACAGATATTGAACAAATTGAAGAAGTTACAAACTTT<br>GATTGGACAAACAAAACAAATCTTTTAGGTGTTTTTAAAAACACCTATGTATCTGTATTCGGATACAAC<br>ATGGGTAATCATGGTGACTTCTTATCAAGCGTTGAATATCCAAGCTTTAAATCTGCTAATACAAATTCA<br>AATTTAACCTTAAGCAAAGAATTTAAAGAAATGGATATCAAAGATTATAAATATACATGCGGTGTCGAA<br>AAAGAGTGGCAATCCTCCAGTGTTCTTTACATAGAGTATGCTAAATACTCAAAGTCTTCCAAAGAATTT<br>GAGGAAATCGGAGGGGGAAGCCTCTCCTGCTCTGAGTGGGGAGCTGCTGATGAAGGATATGCCACTGCC<br>AATTTAGTATCATCCTTTAAGAATTTCTTTGATCGGGATGAATGGTATACTGTGTATACCTGCGGCAAC<br>CGCAACACCTCTAAGGTTTCAAGTTACAAAAAAGAAGAGGTGGCGTAA-3’ |
| AI7 | 5'-ATGAATTTTCCGAGAGCATCAAGACTGATGCAAGCAGCAGTTCTGGGCGGCCTGATGGCAGTTTCA<br>GCAGCAGCAACAGCAAAGCAGGATAACGTCAAAGCGAACAGTGACCCTTACGCTGACCAAATTGAAGAA<br>AACAGCACGCCGTTTACCCTTTATGCGTTTACAGTTTTCACAAAGAGCAGCTTCGGAGCGGATGCCATG<br>TACTATCCGACAAACAATGGAGGCACCTATGGCGGGATTGCCATTATTCCCGGCTATACAGCTAAGCTG<br>GCTTCTTTGAAATGGTGGAAAGATTATGAAGTTGCGCATCAATATGTCTGCTTAACGATTGACTTAAAC<br>AGCCTGATGGATGAAACGAGCCAGCTCGGCACACAAAAGTGTGCGGTGCTTCGGGAGGTTGCCTCCATG<br>AAGGGAAAAGTATCTACCCCACTATTCGGCACCATCAACAAAAATAAGTGGGGAATTGCAAGCTGGAAA<br>GTGTCTAACAAAGGTGACTTCTTAACGGCTGTCCATAAAAACAGCTTTAAAACAGGCTGGAATCCGGCT<br>TTCTGGGATCAAAACTCTATCTATGACGGAAAAACATTCCCTTGGTTCCTGACATCACTGGAGCACAGC<br>AAAGCGAAGCCGTCCTCAAATCTATTTAATACGGTTTATGACAGCATGAGCAATTCAAGCAAGAAGTTT<br>ATTGAAATCTGGGGAGCGTCGCACAACTGCGCCGAGGTTGGCACTGCTGATGAAGGCTTTATCAACGGG<br>AAATCCGTTTCAGTCTTTAATCGATTCTTCGAACTCGATAAAAAATACCATACATTCCCGTGCAGCACG<br>CCGGTCGATAAACGTGTTGCCAATTATTCTACAAAATACTGCCAGTAA-3’ |
| AI8 | 5'-ATGAATTTTCCGAGAGCATCAAGACTGATGCAAGCAGCAGTTCTGGGCGGCCTGATGGCAGTTTCA<br>GCAGCAGCAACAGCACAAACAAATCCATATGCAAGAGGACCAAATCCAACAGCAGCATCATTAGAAGCA<br>AGCGCAGGACCATTTACAGTAAGATCATTTACAGTAAGCCGTCCAAGCGGATATGGAGCAGGAACAGTT<br>TATTATCCAACAAATGCAGGCGGAACAGTAGGAGCAATTGCAATTGTTCCAGGATATACAGCACGTCAG<br>TCAAGCATTAAATGGTGGGGACCAAGATTAGCATCACACGGATTTGTTGTTATTACAATTGATACAAAC<br>TCAACATTAGATCAACCATCAAGCCGTTCATCACAACAAATGGCTGCATTACGTCAAGTAGCATCATTA<br>AATGGAACAAGCAGCAGTCCAATTTACGGAAAAGTTGATACAGCAAGAATGGGTGTAATGGGCTGGTCA<br>ATGGGCGGAGGCGGATCATTAATTTCAGCAGCAAATAATCCATCATTAAAAGCAGCAGCACCACAAGCA<br>CCATGGGATAGCTCAACAAACTTCTCATCAGTAACAGTTCCAACATTAATTTTCGCTTGTGAAAATGAT<br>AGCATTGCACCAGTTAACTCATCAGCATTACCAATTTATGATAGCATGTCACGTAATGCAAAACAATTC<br>CTTGAAATTAATGGTGGATCACATTCATGTGCAAACAGCGGAAATAGCAATCAAGCATTAATCGGTAAA<br>AAAGGTGTAGCTTGGATGAAACGTTTCATGGATAATGATACACGTTACTCAACATTCGCTTGTGAAAAT<br>CCAAACAGCACACGTGTATCAGATTTCCGTACAGCAAACTGCAGCTAA-3’ |
| AI9 | 5'-ATGAATTTTCCGAGAGCATCAAGACTGATGCAAGCAGCAGTTCTGGGCGGCCTGATGGCAGTTTCA<br>GCAGCAGCAACAGCACAAACAAATCCGTATGCTCGTGGTCCTAATCCTACTGCAGCTTCATTAGAAGCT<br>AGTGCTGGACCATTTACAGTAAGATCTTTTACTGTAAGTCGTCCAAGTGGATATGGTGCTGGAACAGTA |

| | |
|---|---|
| | TATTATCCAACTAATGCTGGAGGTACAGTAGGAGCAATTGCTATTGTTCCTGGATATACTGCAAGACAA<br>TCTAGTATTAAATGGTGGGGACCACGTTTAGCATCTCATGGATTTGTTGTAATTACTATTGATACTAAT<br>TCAACATTAGATCAGCCATCAAGCCGTTCATCTCAACAAATGGCTGCTTTAAGACAAGTAGCTTCTTTA<br>AATGGAACTAGTAGTAGTCCTATTTATGGAAAAGTTGATACTGCTAGAATGGGAGTTATGGGTTGGTCT<br>ATGGGTGGTGGTGGATCTTTAATATCAGCTGCTAATAATCCTTCATTAAAAGCAGCTGCACCACAAGCT<br>CCATGGGATAGCTCAACTAATTTCTCTTCTGTTACAGTACCAACTTTAATTTTTGCATGTGAAAATGAT<br>AGTATTGCACCAGTAAATTCATCTGCATTACCAATATATGATAGTATGTCTAGAAATGCTAAACAATTT<br>TTAGAAATTAATGGTGGATCACATTCATGTGCAAATAGTGGAAATAGTAATCAAGCATTAATTGGTAAA<br>AAAGGTGTTGCTTGGATGAAAAGATTTATGGATAATGATACACGTTATTCTACATTTGCATGTGAAAAT<br>CCAAATAGTACTAGAGTTTCTGATTTTAGAACTGCTAATTGTAATTAA-3’ |
| AI10 | 5'-ATGAATTTTCCGAGAGCATCAAGACTGATGCAAGCAGCAGTTCTGGGCGGCCTGATGGCAGTTTCA<br>GCAGCAGCAACAGCACAAACCAATTCGCACGTTACCGGTCCATCTGCGAAAGCCGCTTCGCTTTCGGCG<br>GCTGCTGGTTCATTTAAGGTTGTTTCGTTTCATGTGTCAAAACCATACGGAAAAGGCGCAGGAACAGTT<br>TATTACCCAGTCAATGACGGGGGAGCCGTCGGAGCGATTGCAGTCAGGCCTGATTACAGTGTCCGCGGC<br>ACAAGTATTAAATGGTGGGGTCCGCGGCTTGCGTCGCACGGATTTGTCGTCATCACTGTTGATGAAAAC<br>AGTACATTGAATCAGCCGTCCAGCCGATCATCACAGCAGATTGCCGCATTGAAGCAGGTGCCATCCCTG<br>AACGGAACAGGATCAAGCCCGATTTACGGAAAGGTTGATACATCCAGAATGGGCATAATGGGCTGGTCG<br>ATGGGCGGAGGCGGATCCATCATCAGCGCCGCAAACAATCCTGGGCTTAAGGCCGCGGCACCGCAGGCT<br>CCATATGATGCTTCCGTCAGTTTTTCAAAAGTAACCGTGCCGACGCTCATTTTTGCCTGTGAAAATGAT<br>CAAATTGCACCGGTTAACTCGTCAGCTTTACCGATTTATGACGGAATCAGCAAAAATGCTGCCCAATAT<br>CTTGAGATTAACGGGGGATCTCATTCTTGCGCGAAAAGCGGCAATGCCAATCAAGCGTTGCTCGGTCCA<br>AAAGGTGTGGAATGGATGAAACAGTTTATGGATCATGATACAAGATATTCGACATTTGCTTGTCAAAAC<br>CCAAATAGCACAAGGGTTTCTGATTTTAGAACAGCTCATTGCCAGTAA-3’ |
| AI11 | 5'-ATGAATTTTCCGAGAGCATCAAGACTGATGCAAGCAGCAGTTCTGGGCGGCCTGATGGCAGTTTCA<br>GCAGCAGCAACAGCACAAACCAATTCGCACGCATACGGTAAAGATCCGACAGGAAAATCGATTGAAGCT<br>TCTGTCGGACCATTTAAAGTGCTGTCTTATACGGTGAAAAAACCGTCTGGATATGGAAAAGGAACCGTT<br>TACTATCCAGTCAATGCTGACGGAACAGTCGGAGCGATTGTGATCGCTCCCGGCTATACGGCAAGGGAA<br>GCCGATGTAAAATGGTGGGGTTCCCGTTTAGCCTCACATGGCTATATGGTTGTCGATATAGACCCGAAC<br>TCTACCCTTGACAAAAAAGAGTCAAGATCATCACAGCAAATGGCGGCATTGAAGCAGGTGAAATCCATT<br>AAAGGGACGACCACCAATCCGATCTACGGATTGGTTGACACAAACCGTCTGGCCATGATGGGCTGGTCA<br>ATGGGCGGAGGCGGATCCATTTTGATGGCTGCAGACAATCCTTCTATCAAAGTGGCCGCTCCGCAGGCT<br>CCATTTGACGGAACAGGGAATTTTAGTAAAATAACCGTTCCAACGCTCATTTTTGCTTGTGAAAATGAT<br>CAAATTGCACCAACTAATTCATCGGCATTCAATATTTATAATGCTTTTTCAAAAGATCCTTCCAAATTT<br>GTGGAAATTAACGGAGGATCTCATTCTTGCAGTAATTCTGGAAATAGTAATCAAAATTTAATTGGTAAA<br>AAAGGTGTAGAATGGATGAAACAATTTATGAATAATGATACAAGATATTCAACTTTTGCATGTAAAAAT<br>CCAAATAGTACAAAGGTTTCAAGTTTTAGAACTGCAAATTACAGTTAA-3’ |
| AI12 | 5'-ATGAATTTTCCGAGAGCATCAAGACTGATGCAAGCAGCAGTTCTGGGCGGCCTGATGGCAGTTTCA<br>GCAGCAGCAACAGCACAAACAAATCCATATGCAAGAGGACCAAATCCAACAGCAGCATCATTAGAAGCA<br>AGCGCAGGACCATTTACAGTAAGATCATTTACAGTAAGCCGTCCAAGCGGATACGGAGCAGGAACAGTT<br>TATTATCCAACAAATGCAGGCGGAACAGTAGGCGCAATCGCAATTGTTCCTGGATACACAGCAAGACAA<br>TCAAGCATCAAATGGTGGGGACCACGTTTAGCTTCACACGGCTTTGTTGTAATCACAATTGATACAAAC<br>TCAACACTTGACCAACCATCAAGCCGTTCATCACAACAAATGGCTGCATTACGTCAAGTTGCATCATTA<br>AACGGAACAAGCAGCAGCCCAATCTACGGAAAAGTTGACACTGCACGTATGGGTGTAATGGGTTGGTCA<br>ATGGGCGGCGGCGGCTCATTAATCTCTGCAGCAAACAACCCATCATTAAAAGCAGCTGCTCCACAAGCT<br>CCATGGGACAGCTCAACAAACTTCTCATCAGTAACAGTTCCAACATTAATCTTCGCATGTGAAAACGAC<br>AGCATCGCTCCAGTTAACTCATCTGCACTTCCAATCTACGACAGCATGTCACGTAACGCAAAACAATTC<br>TTAGAAATCAACGGCGGATCACACTCATGCGCAAACAGCGGAAACAGCAACCAAGCATTAATCGGTAAA<br>AAAGGTGTTGCTTGGATGAAACGCTTCATGGACAACGACACACGCTACTCAACATTCGCTTGCGAAAAC<br>CCAAACAGCACACGTGTATCTGACTTCCGTACAGCAAACTGCAGCTAA -3’ |

### Supplementary Figures

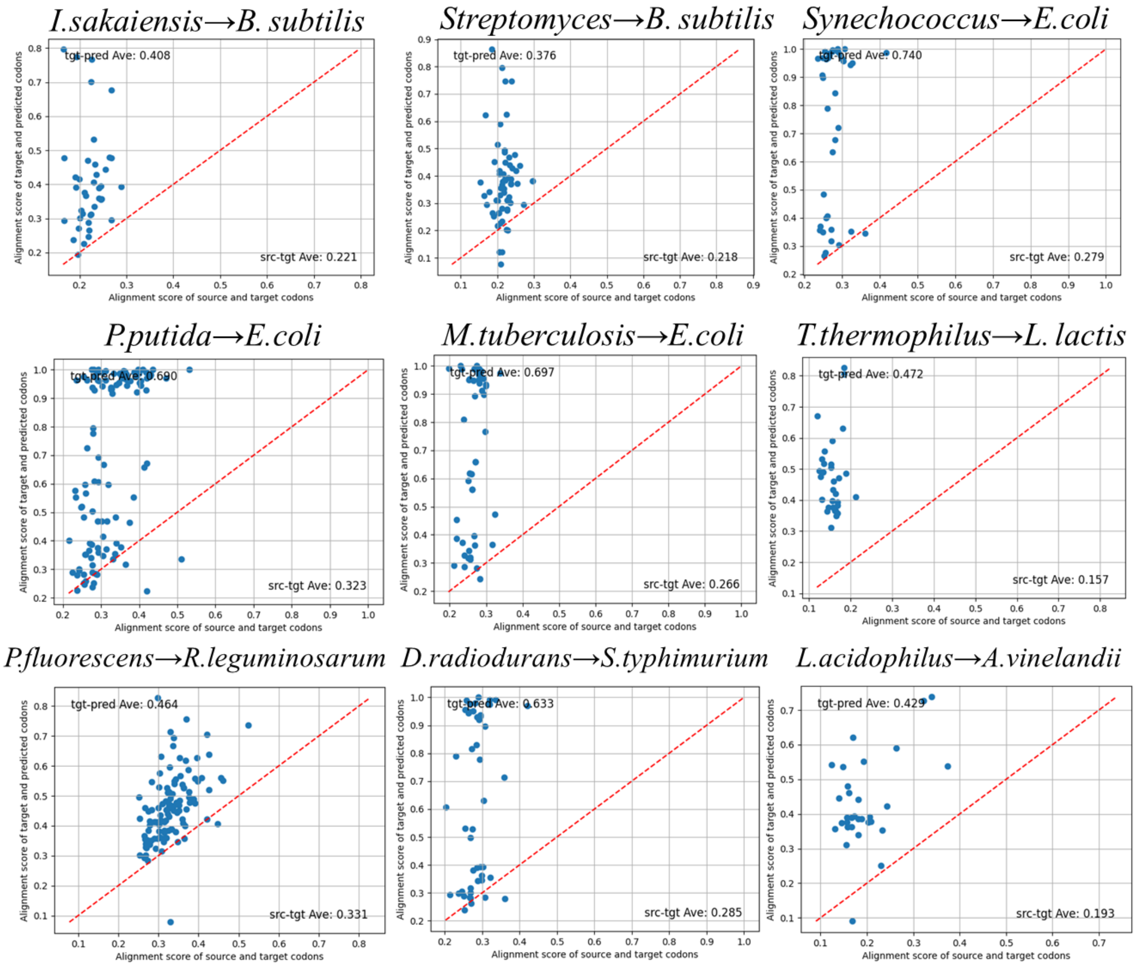


**Supplementary Figure 1.** Codon-level conversion accuracy across diverse bacterial species pairs.

Each scatter plot compares similarity scores between target sequences and AI-generated sequences (y-axis) against similarity scores between target sequences and original source sequences (x-axis) for each bacterial species pair. Individual points represent orthologous gene pairs. Points located above the diagonal red line indicate successful conversions, meaning the AI-generated sequence is more similar to the target sequence than the original source sequence was. The average similarity scores for each comparison are displayed within each plot: "tgt-pred Ave" denotes the average similarity between the target and AI-predicted sequences, whereas "src-tgt Ave" indicates the average similarity between source and target sequences.

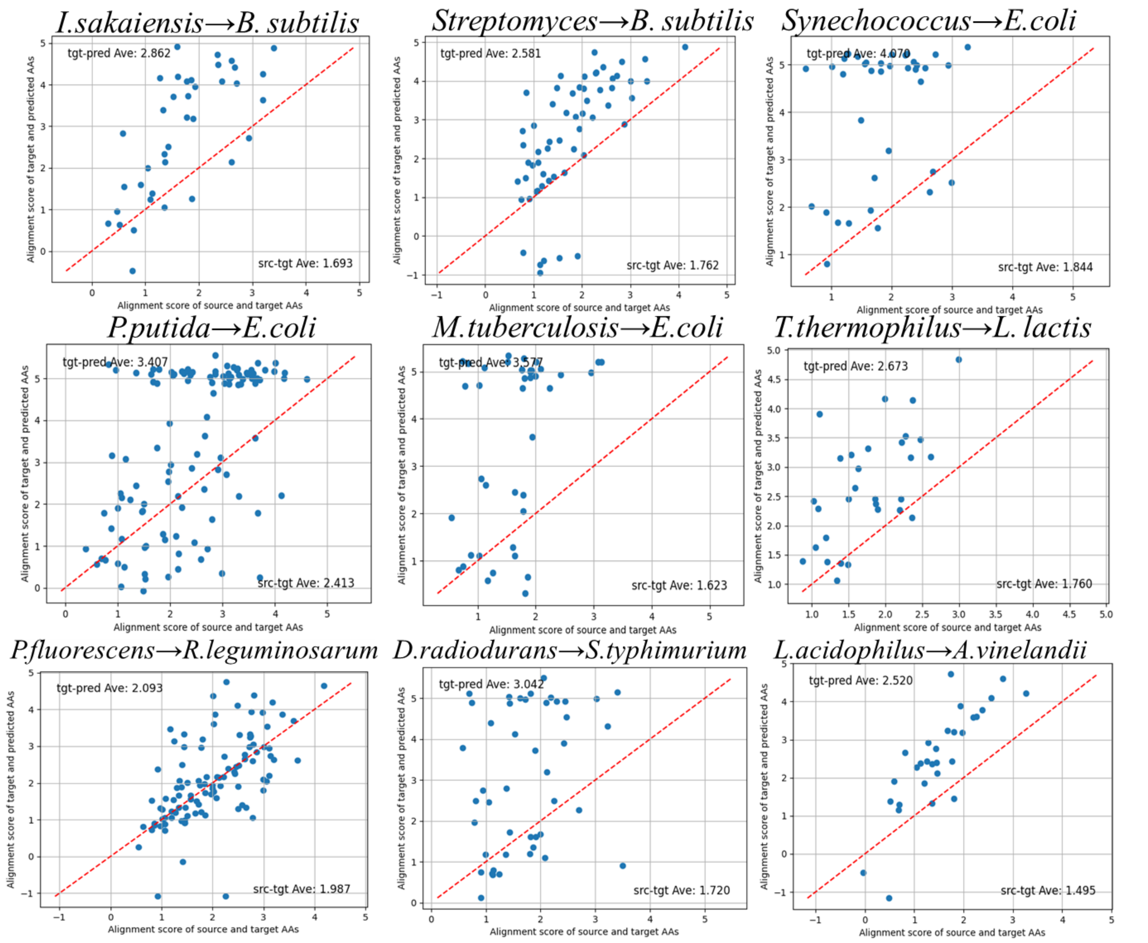


**Supplementary Figure 2.:** Amino acid-level conversion accuracy using BLOSUM score analysis.

To evaluate the functional conservation of generated sequences beyond simple sequence identity, we analyzed amino acid substitution patterns using BLOSUM scores. Each scatter plot shows BLOSUM-based alignment scores between target and generated sequences (y-axis) versus scores between target and source sequences (x-axis). Higher BLOSUM scores indicate more conservative substitutions that preserve protein function.

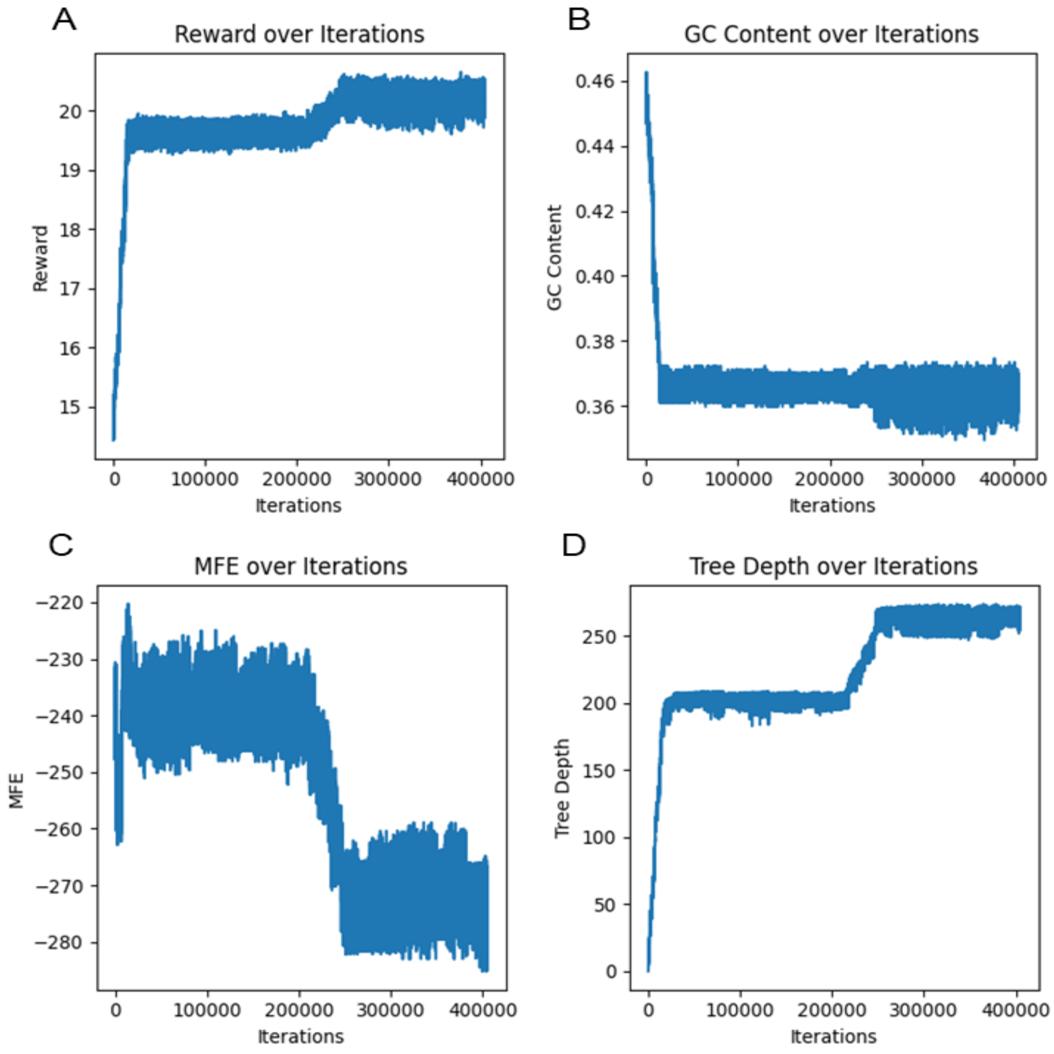


**Supplementary Figure 3.** MCTS optimization trajectories for PETase sequence design.

The figure shows the progression of four key metrics during Monte Carlo Tree Search optimization: (A) Reward value, (B) GC content, (C) Minimum Free Energy (MFE), and (D) Tree depth over 400,000 iterations. The two-phase optimization pattern is evident, with rapid initial improvement followed by more gradual refinement. The algorithm successfully navigated the trade-off between reducing GC content (target: 0.36) and maintaining RNA secondary structure stability (indicated by lower MFE values).

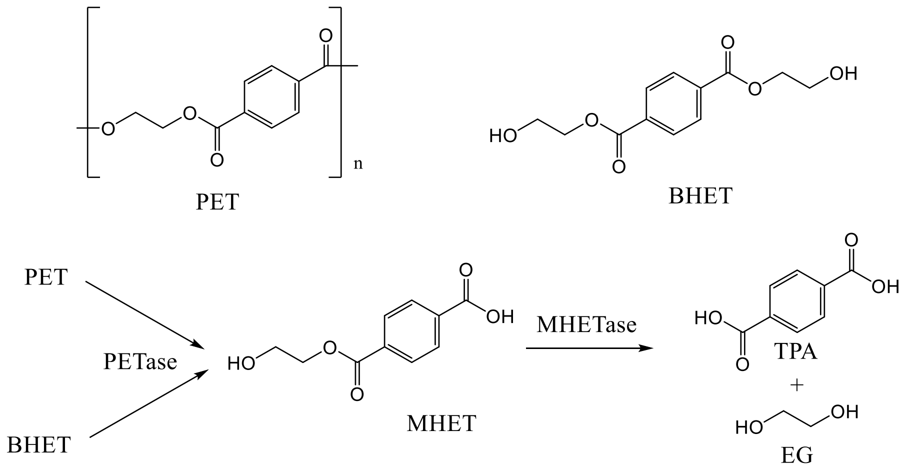


**Supplementary Figure 4.** PET degrading pathway

PETase hydrolyzes PET (polyethylene terephthalate) and BHET (bis(2-hydroxyethyl) terephthalate) into MHET (mono(2-hydroxyethyl) terephthalate). The produced MHET is further hydrolyzed by MHETase into TPA (terephthalic acid) and EG (ethylene glycol).

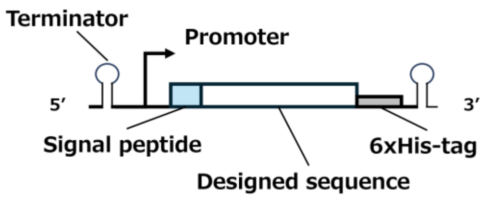


**Supplementary Figure 5.** Sequence design

Gene expression is driven by the $P_{xyl}$ promoter and induced by xylose addition. Terminator sequences are incorporated to prevent transcriptional read-through from upstream and downstream regions. All constructs feature an N-terminal secretion signal peptide for Sec-pathway export and a C-terminal 6×His tag for detection.

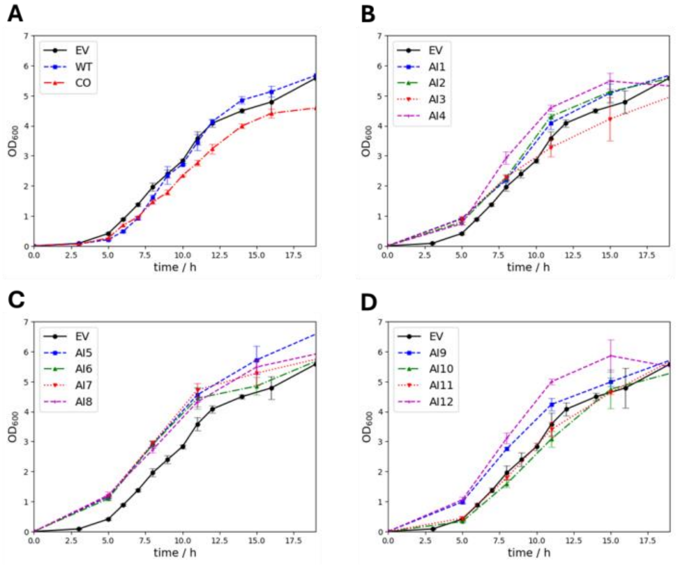


**Supplementary Figure 6.** Growth curve

All recombinant strains were cultured in LB medium supplemented with 25 µg/mL kanamycin and 1% xylose at 30 °C with shaking at 200 rpm, and cell growth was monitored by measuring the optical density at 600 nm ($OD_{600}$). WT and CO (A), AI1~4 (B), AI5~8 (C) and AI9~12 (D).

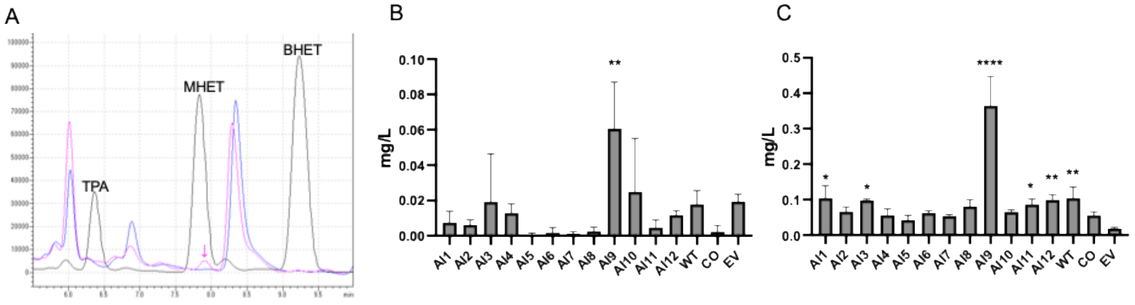


**Supplementary Figure 7.** HPLC analysis of PET degradation by AI-designed PETase variants.

(A) HPLC analysis of Standard solution of TPA, MHET and BHET (black line), culture medium of AI9 and PET film (pink line) and culture medium of AI9 without PET film (blue line). HPLC analysis of MHET in culture medium of engineered strains with PET film for 2days (B) and 7 days (C). All experiments were performed at least in triplicate. Error bars indicate standard deviation. *p < 0.05, **p < 0.01, ****p < 0.0001 (Ordinary one-way ANOVA).

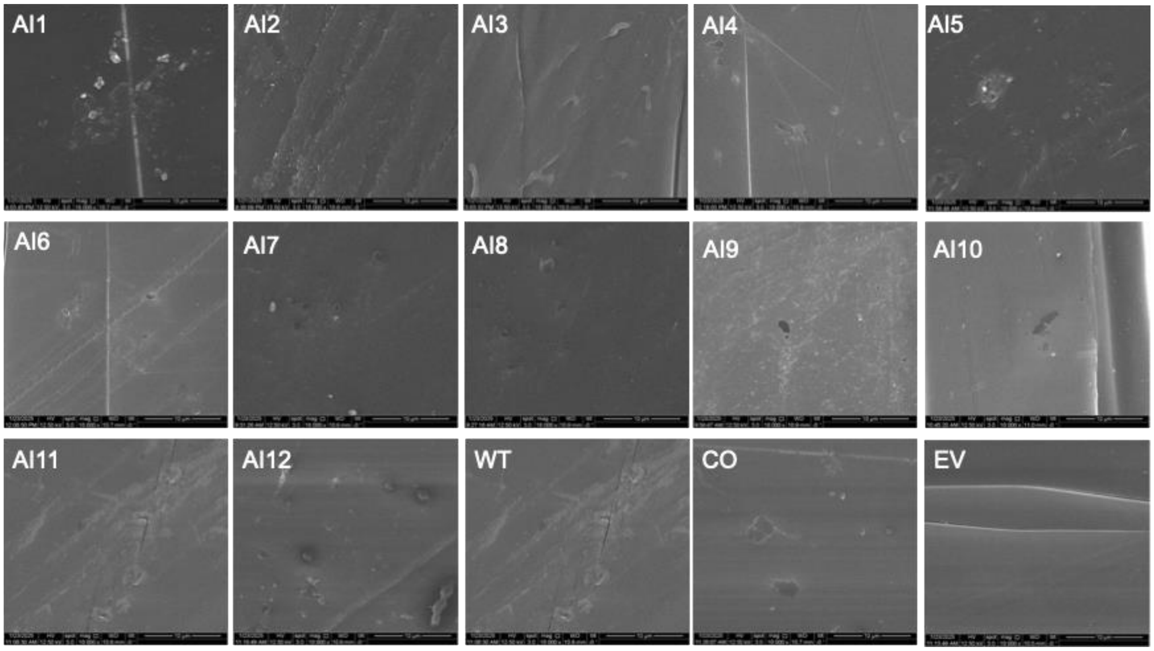


**Supplementary Figure 8.** Degradation of PET film treated with engineered strains for 7 days, as visualized by SEM.

Supplementary References

[S1] Noskov, V. N. et al., Assembly of large, high G+C bacterial DNA fragments in yeast, *ACS Synth. Biol.*, vol. 1, no. 7, pp. 267–273, Jul. 2012.

[S2] Lin, Y. et al., Integrating Reinforcement Learning and Monte Carlo Tree Search for enhanced neoantigen vaccine design, *Brief. Bioinform.*, vol. 25, no. 3, p. bbae247, Mar. 2024.

[S3] Leppek, K. et al., Combinatorial optimization of mRNA structure, stability, and translation for RNA-based therapeutics, *Nat. Commun.*, vol. 13, no. 1, p. 1536, Mar. 2022.

[S4] Zhao, X. et al., High copy number and highly stable Escherichia coli-Bacillus subtilis shuttle plasmids based on pWB980, *Microb. Cell Fact.*, 19, 25 2020.

[S5] Uozumi, T. et al., Restriction and modification in Bacillus species: genetic transformation of bacteria with DNA from different species, part I, *Mol. Gen. Genet.*, 152, 65–69 1977.
